# Deuterium Metabolic Imaging Phenotypes Mouse Glioblastoma Heterogeneity Through Glucose Turnover Kinetics

**DOI:** 10.1101/2024.06.23.600246

**Authors:** Rui V. Simões, Rafael N. Henriques, Jonas L Olesen, Beatriz M. Cardoso, Francisca F. Fernandes, Mariana A.V. Monteiro, Sune N Jespersen, Tânia Carvalho, Noam Shemesh

## Abstract

Glioblastomas are aggressive brain tumors with dismal prognosis. One of the main bottlenecks for developing more effective therapies for glioblastoma stems from their histologic and molecular heterogeneity, leading to distinct tumor microenvironments and disease phenotypes. Effectively characterizing these features would improve the clinical management of glioblastoma. Glucose flux rates through glycolysis and mitochondrial oxidation have been recently shown to quantitatively depict glioblastoma proliferation in mouse models (GL261 and CT2A tumors) using dynamic glucose-enhanced (DGE) deuterium spectroscopy. However, the spatial features of tumor microenvironment phenotypes remain hitherto unresolved. Here, we develop a DGE Deuterium Metabolic Imaging (DMI) approach for profiling tumor microenvironments through glucose conversion kinetics. Using a multimodal combination of tumor mouse models, novel strategies for spectroscopic imaging and noise attenuation, and histopathological correlations, we show that tumor lactate turnover mirrors phenotype differences between GL261 and CT2A mouse glioblastoma, whereas recycling of the peritumoral glutamate-glutamine pool is a potential marker of invasion capacity in pooled cohorts, linked to secondary brain lesions. These findings were validated by histopathological characterization of each tumor, including cell density and proliferation, peritumoral invasion and distant migration, and immune cell infiltration. Our study bodes well for precision neuro-oncology, highlighting the importance of mapping glucose flux rates to better understand the metabolic heterogeneity of glioblastoma and its links to disease phenotypes.

## 1. Introduction

Glioblastoma (glioma grade 4 or GBM) are the most aggressive primary brain tumors in adults. The dismal prognosis of such heterogeneous tumors is mostly attributed to recurrence, associated with limited response to treatment and an infiltrative pattern that prevents full surgical resection [1]. Glioblastoma heterogeneity is reflected in the tumor microenvironment, where glioma cells constantly adapt to their evolving microhabitats, with different biophysical characteristics, progression stages, and therapy resistance [2]. To sustain active proliferation, cancer cells exchange metabolic intermediates with their microenvironment [3] and undergo metabolic reprogramming [4], relying heavily on aerobic glycolysis – upregulation of glucose uptake concomitant with lactate synthesis, leading to acidification of the tumor microenvironment. While this so-called Warburg effect [5] favors e.g. invasion [6], metabolic plasticity [7, 8] is becoming increasingly associated with malignant phenotypes [9]. Namely, mitochondrial oxidation (e.g. glucose metabolism through the tricarboxylic acid cycle, TCA) is linked with microenvironment adaptation and tumor progression [10].

The ability to use both glycolysis and mitochondrial oxidation pathways is a critical feature of GBM, which has been demonstrated from preclinical models to patients [11–13]. More recently, specific dependencies/proclivities towards those metabolic pathways are beginning to reveal GBM subtypes with prognostic value in human cell lines and patient-derived cells [14–16]. Importantly, the latest WHO classification of central nervous system tumors now distinguishes two metabolic phenotypes of adult GBM based on molecular assessment of a specific TCA cycle mutation (isocitrate dehydrogenase, IDH), namely into grade 2-4 gliomas (IDH-mut) and grade 4 GBM (IDH-wt) [17]. The prognostic value of GBM metabolic phenotypes clearly calls for non-invasive imaging methodologies capable of resolving the different subtypes, both for diagnosis and for treatment response monitoring. However, such methods are scarce.

Deuterium metabolic imaging (DMI) has been proposed for mapping active metabolism *de novo* in several tumor models [18–24]. While this has also been demonstrated in GBM patients, with an extensive rationale of the technique and its clinical translation [18], and more recently in mouse models of patient-derived GBM subtypes [25], mapping glucose metabolic fluxes remains unaddressed in these tumors due to the poor temporal resolution of DMI; particularly for glucose mitochondrial oxidation. Leveraging the benefits and risks of denoising methods for MR spectroscopy [26–28], we recently combined Deuterium Magnetic Resonance Spectroscopy (^2^H-MRS) [29] with Marcheku-Pastur Principal Component Analysis (MP-PCA) denoising [30] to propose Dynamic Glucose-Enhanced (DGE) ^2^H-MRS [31], demonstrating its ability to quantify glucose fluxes through glycolysis and mitochondrial oxidation pathways *in vivo* in mouse GBM, which in turn revealed their proliferation status.

Here, we develop and apply a novel rapid DGE-DMI method to spatially resolve glucose metabolic flux rates in mouse GBM and reach a temporal resolution compatible with its kinetic modeling. For this, we adapt two advances of PCA denoising – tensor MPPCA [32, 33] and threshold PCA denoising [34] – and apply it for regional metabolic assessment of mouse GBM. First, we validated our novel approach *in vivo* for its ability to map glucose fluxes through glycolysis and mitochondrial oxidation in mouse GBM. Then, we investigate the potential of our new approach for depicting histopathologic differences in two mouse models of glioblastoma, including microglia/macrophage infiltration, tumor cell proliferation, peritumoral invasion and migration. For this we used the same allograft mouse models of GBM, induced with CT2A and GL261 cell lines [35–39], but at more advanced stages of progression [31]. Since DMI is already performed in humans, including in glioblastoma patients [18], DGE-DMI could be relevant to improve the metabolic mapping of the disease.

## 3. Results

### MRI assessment of mouse GBM

Multi-parametric MRI provided a detailed characterization of each cohort at endpoint. Volumetric T2-weighted MRI indicated consistent tumor sizes across CT2A and GL261 cohorts (58.5±7.2 mm^3^). GL261 tumors were studied sooner after induction (17±0 vs 30±5 days post-injection, p=0.032), explaining the lower animal weights in this cohort (22.4±0.6 vs 25.7±0.9 g, p=0.017). DCE T1-weighted MRI indicated higher vascular permeability (0.85±0.11 vs 0.43±0.05 ·10^-2^/min, p=0.012) and a tendency for larger extracellular volume fractions (0.26±0.03 vs 0.18±0.02, p=0.056) in the GL261 tumors compared to CT2A. However, DCE T1-weighted MRI was carried out only in 80% of the mice due to time restrictions. This information is detailed in **Table S1**, where quantitative assessment of DGE-DMI, DCE-T1 and histologic parameters is displayed for tumor and peritumor border regions (P-Margin), based on ROI analysis.

### DGE-DMI in mouse GBM

Tumor metabolic assessment was performed with DGE-DMI in CT2A vs GL261 cohorts. No differences in RF coil quality or magnetic field homogeneity were detectable between the two cohorts: Q-factor ^2^H, 175±8 vs 176±9 (p=0.8996), respectively; FWHM ^1^H (VOI), 29.2±6.6 vs 26.0±4.3 Hz (p=0.3837), respectively. DGE-DMI was used to map the natural abundance semi-heavy water signal (DHO) as well as the dynamic conversion of deuterium-labelled glucose (Glc) to its downstream products, lactate (Lac) and glutamate-glutamine (Glx) pools, in tumor and peritumor brain regions (**Fig. 1A**). Tensor PCA denoising improved the spectral quality compared to the original data, without any depictable effects in the relative spatial distributions of signal-to-noise-ratio (SNR, **Fig. S1**), leading to a consistent and significant ∼3-fold SNR increase across all the subjects (from 6.4±0.1 before denoising to 20.1±0.4 after denoising, **Table S1**).

**Fig. 1.**
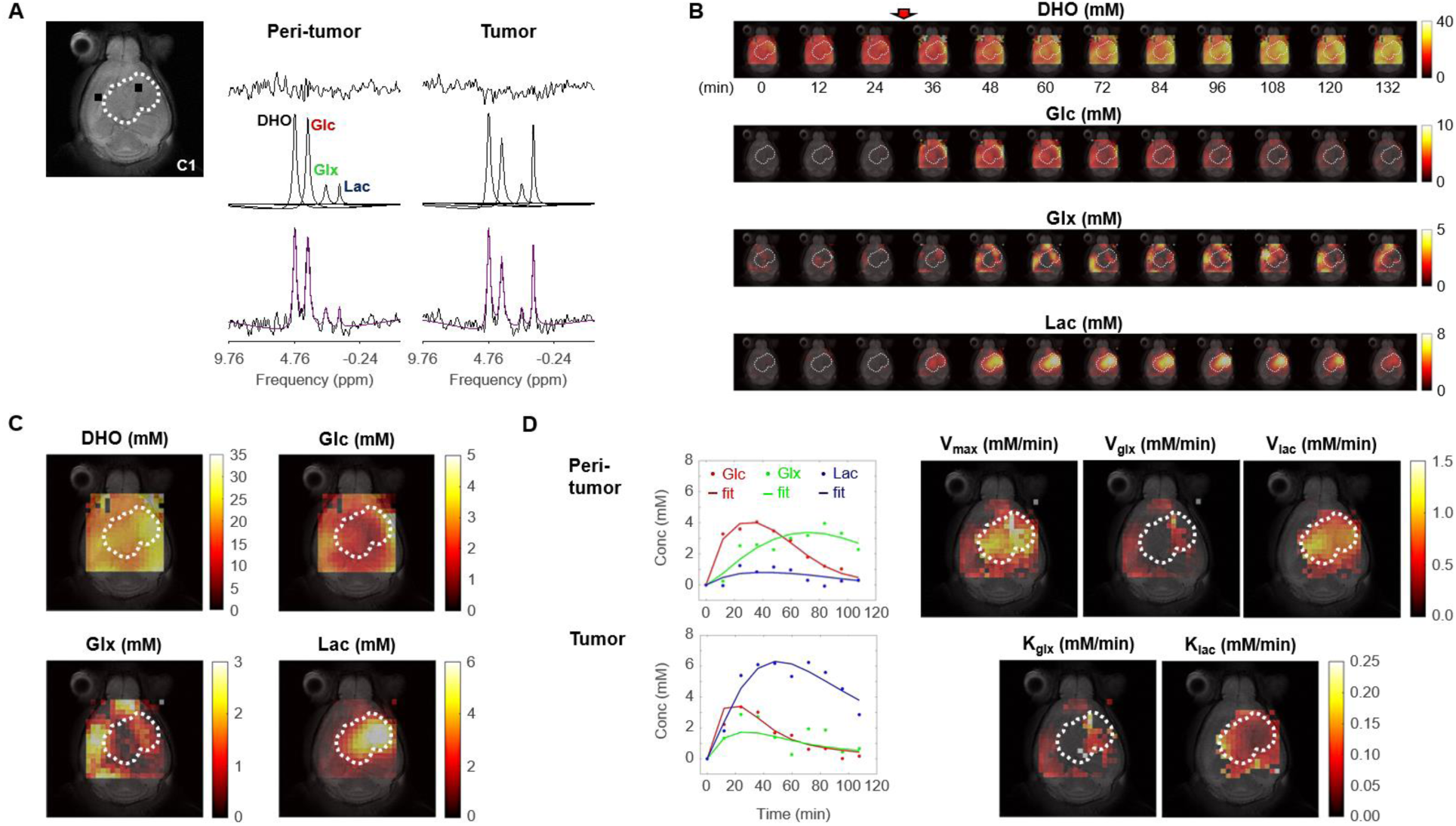
Metabolic concentration and flux maps from DGE-DMI in mouse GBM. Example of a CT2A tumor (C1). **A** T2-weighted reference image (top-left) displaying the tumor region (dashed lines) and representative peritumor and tumor voxels (back dots), and respective spectral quantifications (right-side): bottom, raw spectrum (black) with overlaid estimation (purple); center, individual components for each metabolite peak (black - semi-heavy water, DHO (black); deuterated glucose, Glc (red); and glucose-derived glutamate-glutamine and lactate, Glx (green) and Lac (blue)); top, residual. **B** Time-course *de novo* concentration maps for each metabolite (mM) following Glc i.v. injection (red arrow). **C** Average concentration maps for each metabolite after Glc injection. **D** Time-course concentration plots for each metabolite (dots) and respective kinetic fitting (straight lines), displayed for the peritumor and tumor voxels shown in A (same color codes) and applied to all the voxels to generate glucose flux maps: maximum consumption rate (V_max_); and respective individual rates for lactate synthesis (V_lac_) and elimination (k_lac_), and glutamate-glutamine synthesis (V_glx_) and elimination (k_glx_).

Spectral quantification of DGE-DMI data in each voxel and time point rendered time-course *de novo* concentration maps for each metabolite (DHO, Glc, Glx, and Lac), in both GBM cohorts (**Fig 1B**). Voxel-wise averaging of DGE-DMI time-course data after Glc injection generated average metabolic concentration maps for each tumor (**Fig. 1C**). Thus, Lac concentration was visually higher in the tumor regions, due to enhanced glycolysis; whereas Glx was more apparent in the adjacent non/peritumoral areas, consistent with a more prevalent oxidative metabolism in the normal brain. Kinetic fitting of DGE-DMI time-course concentration maps rendered glucose flux maps, namely its maximum consumption rate (V_max_) and flux rates through glycolysis (V_lac_ and k_lac_) and mitochondrial oxidation (V_glx_ and k_glx_) (**Fig 1D**). Both cohorts displayed higher glycolytic metabolism in the tumors and more pronounced glucose oxidation in non-tumor regions, aligned with average concentration maps.

### Histopathology assessment of GBM cohort differences

Histopathological analysis consisted of screening the CT2A and GL261 brain tumors for morphological features, including qualitative assessment of cell density, hemorrhage, tumor vessels, necrosis, quantification of peripheral infiltration and quantification of tumor proliferation index, while blinded to the in vivo MRI/MRS data – **Table S2**. Thus, tumors were scored individually for the following stromal-vascular phenotype, as in [31], where: pattern I corresponds to predominance of small vessels, complete endothelial cell lining and sparse hemorrhages; pattern II to vasodilation and marked multifocal hemorrhages; pattern III to predominance of necrosis of the vascular wall, incomplete endothelial cell lining, vascular leakage, and edematous stroma; and pattern IV to tumors with absence of clear vascular spaces and edematous stroma.

Stromal-vascular phenotypes reflected the more advanced stages of tumor progression in which these tumors were collected, as compared to our previous study [31]. Particularly, CT2A (n=5) presented patterns I to III, whereas all GL261 (n=5) matched pattern IV (**Table S1**). Moreover, the increased infiltrative and migratory characteristics of GL261 compared to CT2A tumors were evident in their irregular tumor borders and higher incidence of secondary brain lesions (**Fig 2A**). These findings collectively suggest a more invasive and aggressive pattern of GL261 tumors, characterized by reduced cell-cell adhesion and enhanced migratory potential compared to CT2A. Such phenotype differences were reflected in the regional infiltration by microglia/macrophages: significantly higher at the CT2A peritumoral margin (P-Margin) compared to GL261, and slightly higher in the tumor region as well (**Fig 2B**). Further quantitative regional analysis of Tumor-to-P-Margin ROI ratios revealed: (i) 47% lower cell density (p=0.004) and 32% higher cell proliferation (p=0.026) in GL261 compared to CT2A (**Fig 2C, Table S3**); and (ii) strong negative correlations in pooled cohorts between microglia/macrophage infiltration and cellularity (R=-0.91, p=<0.001) or cell density (R=-0.77, p=0.016), suggesting more circumscribed tumor growth with higher peripheral/peritumoral infiltration of immune cells.

**Figure 2.**
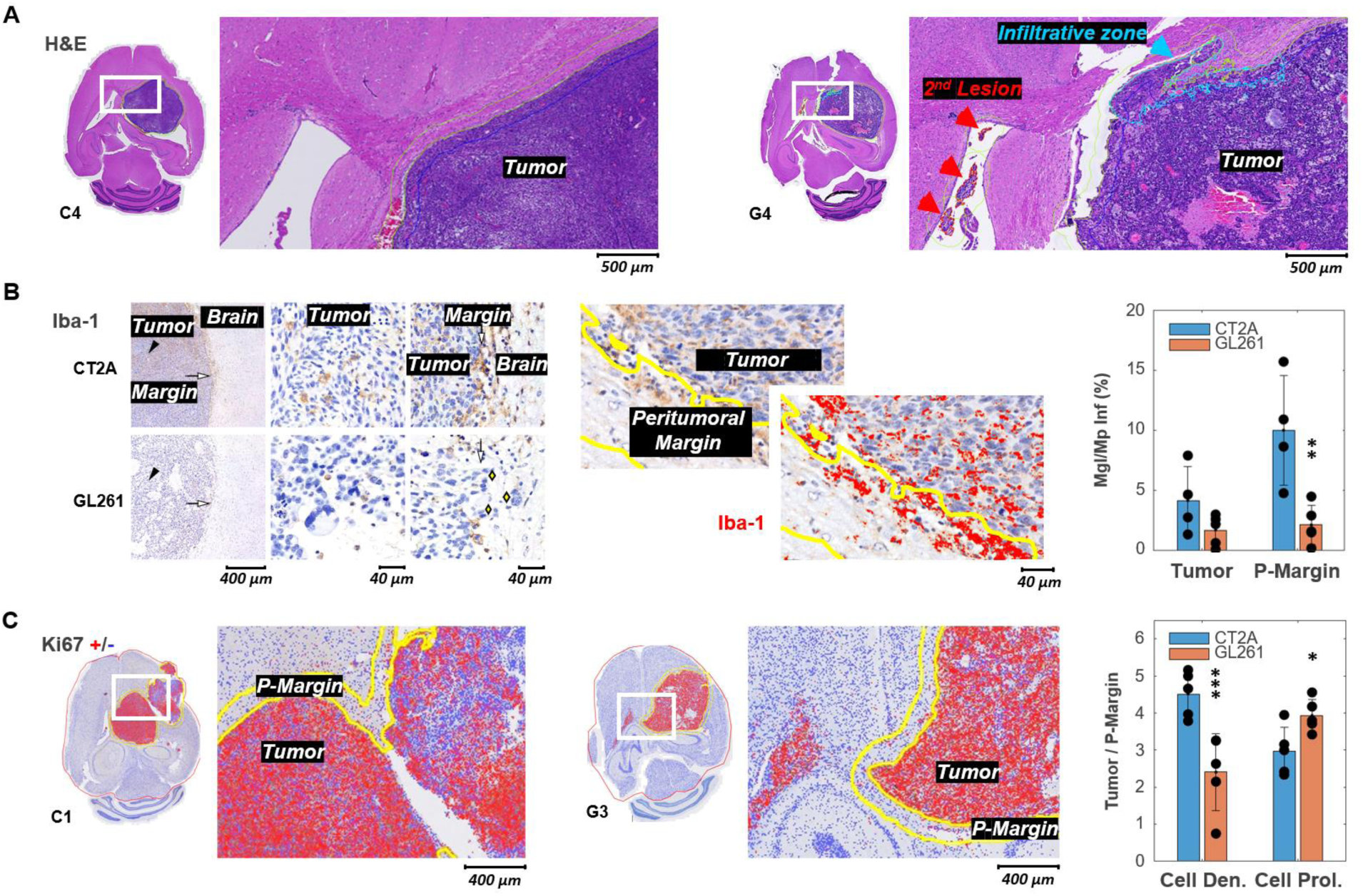
Histopathologic and immunohistochemical assessment in two mouse models of GBM. **A** H&E-stained sections with high magnification to highlight annotations of tumor, infiltrative zones in the tumor margin (blue), and secondary lesion (red), in CT2A and GL261 tumors (subjects C4 and G4, respectively). **B** Iba-1 immunostained sections showing microglia/macrophage (Mgl/Mp) infiltration in CT2A and GL261 tumors: left panels, tumor core (black arrowhead) and tumor margin (white arrow) relative to the adjacent brain parenchyma; middle and right panels, depicting more infiltration by microglial/macrophage in CT2A tumors, also with clearer well-demarcated margin where IBA-1-positive cells are more densely concentrated compared to the more diffuse and irregular infiltration seen in the GL261 model; GL261 show poorly demarcated tumor border where tumor cells infiltrate the brain parenchyma (yellow diamonds); center panels, Iba-1 ROI quantification in tumor and peritumoral margin (P-Margin, yellow lines), and with red mask overlay of Iba-1 positive cells; right panel, quantification of mean Iba-1 positive area in Tumor and P-Margin regions of CT2A and GL261 cohorts – C2 sample excluded due to peritumoral hemorrhage/vascular ectasia, which distorted the peritumoral area and impaired proper assessment of peritumoral infiltration. **C** Ki67 immuno-stained sections with overlaid detection of positive (red) and negative (blue) cells; and high magnification to highlight annotations of tumor and peritumor border (P-Margin, yellow lines), in CT2A and GL261 tumors (subjects C1 and G3, respectively); and GBM cohort differences in tumor/P-Margin ratios of cell density and cell proliferation (dots representative of average values for each subject).. CT2A vs GL261 (ki67, n= 5 vs 5; Iba-1, n= 4 vs 5): * p<0.05; ** p<0.01; *** p<0.001; unpaired *t*-test. Error bars: standard deviation.

Despite the more advanced stages of tumor progression, the results were largely consistent with the marked morphological differences between the two models [31]: CT2A with dense, cohesive and homogeneous cell populations (**Fig 2A**, left-side); GL261 displaying marked heterogeneity, with poorly cohesive areas and more infiltrative growth (**Fig 2A**, right-side). Quantitative assessment (nuclear counts) further confirmed a nearly 2-fold lower cell density of GL261 tumors compared to CT2A (4.9 vs 8.2 ·10^3^ cells/µm^2^, p<0.001) despite their similar proliferation index (**Table S1**); and tumor cell density correlated with cell proliferation, strongly for CT2A (R=0.96, p=0.009) and the same tendency detected for GL261 (R=0.74, p=0.151).

Tumor volume and whole-brain gross assessment of cell density, cell proliferation, and glucose metabolism also revealed strong inter-subject correlations in both cohorts (**Fig. S2**): *de novo* glutamate-glutamine accumulation decreased with tumor size (R _CT2A/ GL261/ pooled_: −0.597/ - 0.753/ −0.455), consistent with its role as marker of oxidative metabolism in the normal brain; lactate synthesis rate increased with cellularity (R _CT2A/ GL261/ pooled_: +0.921/ +0.685/ +0.852), also aligned with enhanced glycolysis in growing tumors; whereas glucose accumulation reflected cell proliferation (R _CT2A/ GL261/ pooled_: +0.469/ +0.528/ +0.440).

### Regional assessment of glucose metabolism in the tumor microenvironment

Initial intra-tumor analysis of DGE-DMI and DCE-T1 maps (pixel-wise correlations in tumor ROIs) indicate stronger correlations between *de novo* lactate accumulation (Lac) and vascular permeability (k^trans^) in both cohorts (R between [+0.306 +0.741]), and extracellular space (v_e_) to some extent (R between [−0.084 +0.804]) – both less apparent without tensor PCA denoising (R between [+0.089 +0.647] and [−0.160 +0.684], respectively) (**Fig S3**). Such accumulation of lactate according to local vascular permeability mostly reflected regional differences in glycolytic fluxes (V_lac_: R between [−0.066 +0.510]), rather than lactate elimination rates (k_lac_: R between [−0.643 +0.460]). No additional correlations were detected.

GL261 tumors accumulated significantly less lactate in the core (1.60±0.25 vs 2.91±0.33 mM: −45%, p=0.013) and peritumor margin regions (0.94±0.09 vs 1.46±0.17 mM: −36%, p=0.025) than CT2A – **Fig 3 A-B**, **Table S1**. Consistently, tumor lactate accumulation correlated with tumor cellularity in pooled cohorts (R=0.74, p=0.014). Then, lower tumor lactate levels were associated with higher lactate elimination rate, k_lac_ (0.11±0.1 vs 0.06±0.01 mM/min: +94%, p=0.006) – **Fig 3B** – which in turn correlated inversely with peritumoral margin infiltration of microglia/macrophages in pooled cohorts (R=-0.73, p=0.027) - **Fig 3-C**. Further analysis of Tumor/P-Margin metabolic ratios (**Table S3**) revealed: (i) +38% glucose (p=0.002) and −17% lactate (p=0.038) concentrations, and +55% higher lactate consumption rate (p=0.040) in the GL261 cohort; and (ii) lactate ratios across those regions reflected the respective cell density ratios in pooled cohorts (R=0.77, p=0.010) – **Fig 3-C**. Finally, lactate elimination rate correlated inversely with “tumor age” (time post-induction) in pooled cohorts (R=-0.66, p=0.039), and more consistently with tumor vascular permeability (k^trans^: R=0.78, p=0.022) (**Fig 3C**), rather than washout rate (k_ep_: R=0.61, p=0.109).

**Figure 3.**
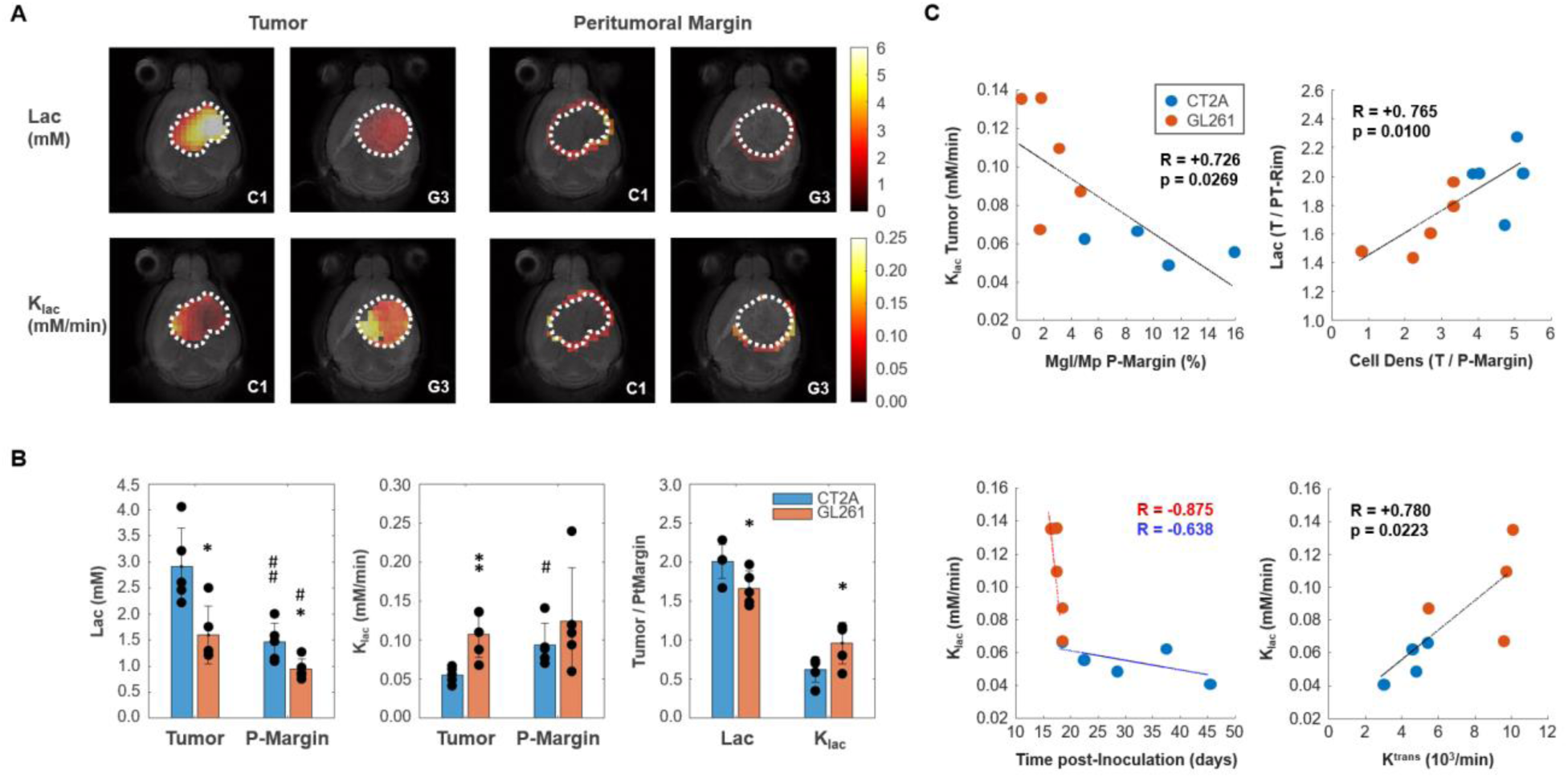
Mouse GBM models with different histopathologic phenotypes underlied by regional differences in lactate metabolism. **A** Metabolic maps of *de novo* lactate accumulation (mM) and respective consumption/elimination rates (mM/min), in tumor and peritumor border regions (P-Margin, delineated by dashed lines) of CT2A and GL261 tumors (subjects C1 and G3, respectively). **B** GBM cohort differences in *de novo* lactate accumulation (Lac) and consumption/elimination rates (k_lac_). **C** Strong linear correlations (indicated by the Person correlation coefficient, R) of: top-left, Tumor lactate consumption/elimination rates with P-Margin infiltration of microglia/macrophages in pooled cohorts; top-right, Tumor-to-P-Margin ratios of lactate accumulation and cell density in pooled cohorts; bottom, lactate consumption/elimination rates with (left-side) time post-tumor inoculation in each cohort, and (right-side) tumor vascular permeability in pooled cohorts. CT2A (n=5) vs GL261 (n=5): * p<0.05; ** p<0.01; unpaired *t*-test. Tumor (n=5, each cohort) vs P-Margin (n=5, each cohort): # p<0.05; ## p<0.01; paired *t*-test. Error bars: standard deviation. Bar plot dots representative of average pixel values for each subject.

### Association between glucose metabolism and peritumoral invasion and migration

Finally, we investigated the association between glucose metabolism and phenotypic features of tumor aggressiveness, namely cell proliferation and tumor cell invasion and migration associated with secondary brain lesions. Only the more infiltrative GL261 cohort displayed inter-subject associations between tumor cell proliferation (Ki67^+^ %) and metabolism, namely inverse correlations with tumor border/peritumoral glucose oxidation rate (V_glx_: R=-0.91, p=0.030) and glucose-derived glutamate-glutamine elimination rate (k_glx_: R=-0.99, p<0.001). Regrouping subjects according to glioma cell invasion and migration concomitant with secondary brain lesions (presence: C1, G3, G4, G5; vs. absence: C2, C3, C4, C5, G1, G2) revealed lower *de novo* glutamate-glutamine levels in peritumor brain regions (Glx: −37%, p=0.013), which were associated with its higher elimination rate (k_glx_: +69%, p=0.012) – **Fig 4**.

**Figure 4.**
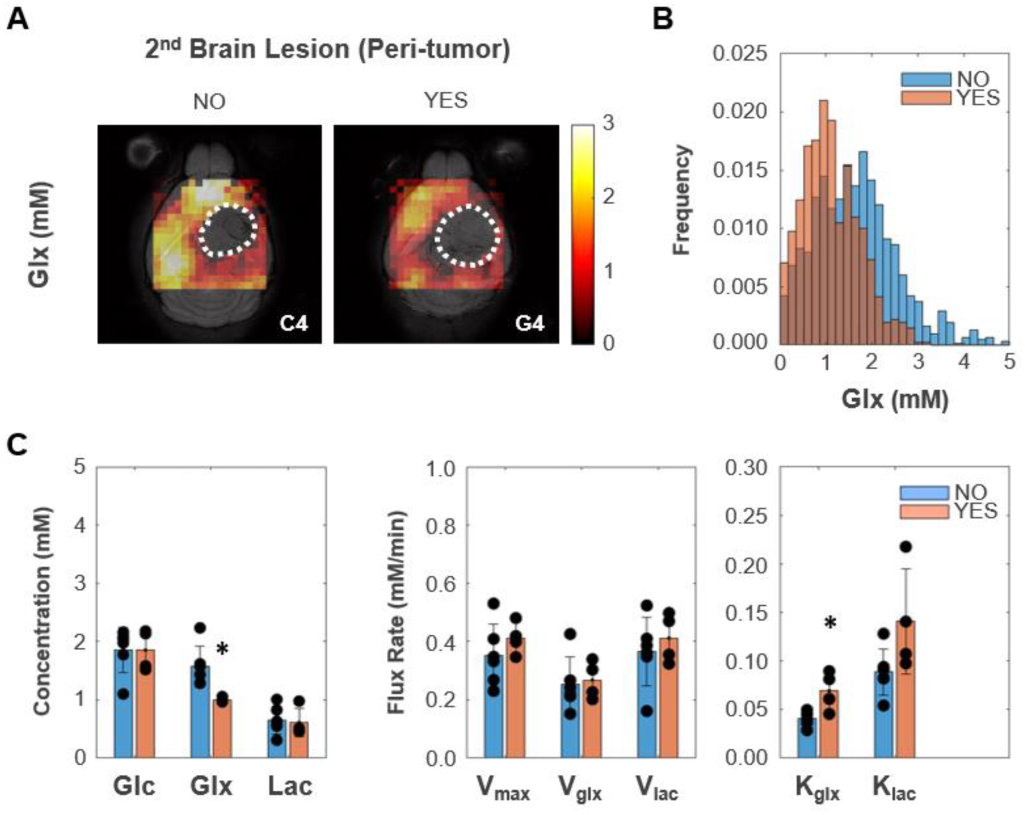
Peritumoral metabolic changes consistent with recycling of the glutamate-glutamine pool mirror GBM infiltration and migration leading to secondary brain lesions. A Metabolic maps (Glx) of peritumoral regions without and with secondary brain lesions (C4 and G4 tumors, respectively). B Histogram distributions of peritumoral Glx accumulation in pooled GL261 and CT2A cohorts displaying secondary brain lesions (n=4) vs without (n=6). C Bar plot comparison of mean values, showing significant decreases in peritumoral glutamate-glutamine accumulation (Glx) and increases in its consumption/elimination (k_glx_) in pooled GL261 and CT2A cohorts displaying secondary brain lesions (n=4; vs n=6 without): * p<0.05; unpaired *t*-test. Error bars: standard deviation. Bar plot dots representative of average pixel values for each subject.

## Discussion

Glioblastomas are aggressive brain tumors with a poor prognosis, largely due to their inter- and intra-tumor heterogeneity and lack of non-invasive methods to assess it. Here we developed and applied a DGE-DMI approach capable of generating metabolic concentration maps and flux rates in two mouse models of glioblastoma, based on unambiguous spectral quantification according to quality criteria. Our results suggest that glycolytic lactate turnover mirrors phenotype differences between the two glioblastoma models, whereas recycling of the glucose-derived glutamate-glutamine pool could underlie glioma cell migration leading to secondary lesions. This information became more readily available when using the tensor PCA method for spectral denoising.

Tensor PCA denoising increased spectral SNR by ∼3-fold, consistently improving spectral quality observed in tumor and peritumoral regions without altering the spatiotemporal profiles of the metabolic concentration maps (**Fig S4**). While this had no apparent effect on metabolic concentration maps (**Figs S5-6**), it significantly improved the kinetic modeling performance (**Fig S7**) and rendered better quality metabolic flux maps in CT2A and GL261 cohorts. Thus, 63% increased pixel detectability enabled capturing more spatial features in the latter without affecting parameter estimates or introducing group differences (**Figs S8-9**).

Gross whole brain analysis revealed strong inter-subject correlations in both cohorts, such as higher lactate synthesis rate with increasing cellularity – consistent with enhanced glycolysis in growing tumors – whereas intra-tumor pixel-wise analysis suggested lactate accumulation according to local vascular permeability, mostly associated with regional differences in glycolytic fluxes. Such pixel-wise analyses might be misleading since *de novo* lactate diffuses quickly within tumor extracellular spaces and peritumoral regions [40], with spatiotemporal dynamics not fully captured by DGE-DMI. Namely, water diffusion in GL261 tumors *in vivo* (apparent diffusion coefficient ∼10^-3^ mm^2^/s [41, 42]) extends beyond the in-plane voxel area (0.56×0.56 = 0.31 mm^2^) during each time frame (12 min). Thus, we focused instead on inter-tumor ROI analysis of glucose metabolic fluxes, in tumor and peritumoral (border) regions.

Compared to our previous study using the same GBM models [31], larger tumors (59±7 vs 38±3 mm^3^) display more disrupted stromal-vascular phenotypes (H&E scores: CT2A I-III vs I; GL261, IV vs I-IV) and weaker cell-cell interactions (lower cohesiveness) (**Table S2**), associated with lower vascular permeability (k^trans^: 6±1 vs 14±1 10^3^/min) and leading to lower glucose oxidation rates (V_glx_: 0.28±0.06 vs. 0.40±0.08 mM/min), but remarkably similar glycolytic fluxes (V_lac_: 0.59±0.04 vs. 0.55±0.07 mM/min). Thus, glycolysis flux rates are relatively well preserved across GL261 and CT2A mouse GBM models, regardless of tumor volume and vascular permeability.

GL261 tumors were examined earlier after induction than CT2A (17±0 vs. 30±5 days, p = 0.032), displaying similar volumes (57±6 vs. 60±14, p = 0.813) but increased vascular permeability (8.5±1.1 vs 4.3±0.5 10^3^/min: +98%, p=0.001), more disrupted stromal-vascular phenotypes and infiltrative growth (5/5 vs 0/5), consistent with significantly lower tumor cell density (4.9±0.2 vs. 8.2±0.3 10^-3^ cells/µm^2^: −40%, p<0.001) and lower peritumoral rim infiltration of microglia/macrophages (2.1±0.7 vs. 10.0±2.3 %: −77%, p=0.008). Such GBM cohort differences were markedly reflected in their regional lactate metabolism. Thus, GL261 tumors accumulated roughly −40% less lactate in tumor and peritumor border regions, associated with +94% higher lactate elimination rate rather than glycolytic rate differences in tumor regions, as could be assumed solely based on metabolic concentration maps.

Tumor vs peritumor border analyses further suggest that lactate metabolism reflects regional histologic differences: lactate accumulation mirrors cell density gradients between and across the two cohorts; whereas lactate consumption/elimination rate coarsely reflects cohort differences in cell proliferation, and inversely correlates with peritumoral infiltration by microglia/macrophages across both cohorts. This is consistent with GL261’s lower cell density and cohesiveness, more disrupted stromal-vascular phenotypes, and infiltrative growth pattern at the peritumor margin area, where less immune cell infiltration is detected and relatively lower cell division is expected [43]. Altogether, our results suggest increased lactate consumption rate (active recycling) in GL261 tumors with higher vascular permeability, e.g. as a metabolic substrate for oxidative metabolism [44] promoting GBM cell survival and invasion [45], aligned with the higher respiration buffer capacity and more efficient metabolic plasticity of GL261 cells than CT2A [31]. While, lactate shuttling within the tumor microenvironment is also reported in other tumor types, between cancer cells [46] and between cancer and stromal cells [47, 48], it should be noted that oxidative phosphorylation inefficiency has been extensively documented in cancer cells, including GBM [49], largely associated with hypoxic niches and in agreement with our measurements of lower glucose oxidation rate (V_glx_) in tumor vs. peritumoral regions.

The lower glucose oxidation rates measured in this study compared with smaller, better perfused tumors [31], are in good agreement with our previous data indicating quick adaptation of this pathway flux according to oxygen availability in the tumor microenvironment [31]. Under such physiological conditions – underlying more advanced progression stages, reflected in more disrupted stromal-vascular phenotypes – tumor glucose oxidation rate was not associated with cell proliferation index, consistent with previous observations [31]. Instead, tumor cell proliferation was inversely correlated with tumor border/peritumoral glucose oxidation rate and glucose-derived glutamate-glutamine elimination rate in more infiltrative GL261 tumors; but not in CT2A. This observation is consistent to some extent with GL261 cells’ and tumor’s ability to modulate mitochondrial metabolism according to their microenvironment (e.g. oxygen availability [31]), which is likely to occur during their progression from more circumscribed/local cell proliferation towards more disrupted stromal-vascular phenotypes, associated with significantly lower peri-tumoral immune cell infiltration and higher tumor invasion compared to CT2A.

Notably, glucose-derived glutamate-glutamine displayed −37% lower levels and +69% higher elimination rate in peritumor regions of mouse brains bearing secondary GBM lesions (respective primary tumors displaying +146% increased glucose oxidation rate, detectable only with tensor PCA denoising – **Fig S10**). This could be associated with glutamate-glutamine-driven mitochondrial metabolism, through the TCA cycle coupled with oxidative phosphorylation (more prevalent in the normal brain) and/or via substrate level phosphorylation for ATP synthesis – glutaminolysis (as reported in glioma cells, e.g. CT2A [50]). While patient-derived xenografts and *de novo* models would be more suited to recapitulate human GBM heterogeneity and infiltration features, and genetic manipulation of glycolysis and mitochondrial oxidation pathways could be relevant to ascertain DGE-DMI sensitivity for their quantification, our observations are well aligned with the pivotal role of mitochondrial metabolism in cancer cells with higher motile potential, as reported in human GBM [51] and in mouse and human breast cancer cell lines [52, 53]. Particularly, the dynamics of glutamate shuttling underlying neuronal-glioma cell communication and promoting GBM infiltration, are increasingly reported by the emerging field of cancer neuroscience [54]. Therefore, our results suggest that glucose mitochondrial metabolism mirrors GBM progression in mouse GL261 and CT2A models: more prevalent in smaller, well perfused tumors, where glucose oxidation rate correlates with tumor cell proliferation [31]; lower in larger, more poorly perfused tumors, where recycling of the glutamate-glutamine pool may reflect a phenotype associated with secondary brain lesions.

Despite the excellent performance of tensor PCA denoising – 3-fold increase in SNR, approaching the original/raw values obtained previously with single-voxel ^2^H-MRS data (SNR∼20, [31]) – no further improvements in SNR could be achieved by free induction decay (FID) averaging within the tumor ROI (**Fig S11**). Therefore, further DGE-DMI preclinical studies aimed at detecting and quantifying relatively weak signals, such as tumor glutamate-glutamine, and/or increase the nominal spatial resolution to better correlate those metabolic results with histology findings (e.g in the tumor margin), should improve basal SNR with higher magnetic field strengths, more sensitive RF coils, and advanced DMI pulse sequences [55]). In the kinetic model, the extracellular volume fraction was fixed to ensure model stability, as previously demonstrated using the tumor average across all subjects [31]. This approximation may not fully reflect the intra- and inter-tumor heterogeneity of this parameter in both cohorts, and may not be representative of its peritumoral regions. Still, we opted for this approach, rather than pixel-wise adjustments according to DGE-T1 extracellular volume fraction maps, given (i) the relative insensitivity of the model to the actual extracellular volume fraction value used [31], also verified in the present study (**Fig S12**); and particularly, because (ii) we did not have DCE-T1 data for the full cohort, thus it was not feasible to perform individual corrections, which in any case would ultimately be prone to error at tumor periphery/border regions, where exact delimitations are typically debatable. Finally, our results are indicative of higher microglia/macrophage infiltration in CT2A than GL261 tumors, which is inconsistent with another study reporting higher immunogenicity of GL261 tumors than CT2A for microglia and macrophage populations [56]. Such discrepancy could be related to methodologic differences between the two studies, namely the endpoint-guided assessment of tumor growth (bioluminescence vs MRI, more precise volumetric estimations) and tumor stage (GL261 at 23-28 vs 16-18 days post-injection, i.e. less time for immune cell to infiltration in our case), presence/absence of a cell transformation step (GFP-Fluc engineered vs we used original cell lines), or perhaps media conditioning effects during cell culture due to the different formulations used (DMEM vs RPMI).

Our results clearly highlight the importance of mapping pathway fluxes alongside *de novo* concentrations to improve the characterization of the complex and dynamic heterogeneity of GBM metabolism. This may be determinant for the longitudinal assessment of GBM progression, with end-point validation; and/or treatment-response, to help selecting among new therapeutic modalities targeting GBM metabolism [57, 58] or monitoring the efficacy of novel immunotherapy approaches [59] beyond conventional chemoradiotherapy [25]. Importantly, DGE-MRI has already been demonstrated in glioma patients with i.v. administration of glucose using Chemical Exchange Saturation Transfer (glucoCEST) and relaxation-based methods [60, 61], to map the spatiotemporal kinetics of glucose accumulation rather than quantifying its downstream metabolic fluxes through glycolysis and mitochondrial oxidation, as we did. The latter could potentially benefit from an improved kinetic model simultaneously assessing cerebral glucose and oxygen metabolism, as recently demonstrated in the rat brain with a combination of ^2^H and ^17^O MR spectroscopy [62]. Moreover, DMI has been demonstrated on a 9.4T clinical MRI scanner [63], benefiting from the higher sensitivity in the much larger human brain compared to mice: 200 cm^3^ [64] and 415 mm^3^ [65], respectively.

In summary, we report a DGE-DMI method for quantitative mapping of glycolysis and mitochondrial oxidation fluxes in mouse GBM, highlighting its importance for metabolic characterization and potential for *in vivo* GBM phenotyping in different models and progression stages. In large mouse GBM tumors, lactate metabolism underlies model-specific features, consistent with faster turnover in more disrupted stromal-vascular phenotypes and mirroring intra-tumor gradients of cell density and proliferation, whereas recycling of the glutamate-glutamine pool may reflect a phenotype associated with secondary brain lesions. Tensor PCA denoising significantly improved spectral signal-to-noise, which helped reveal such associations between regional glucose metabolism and phenotypic features of intra- and inter-tumor heterogeneity. DGE-DMI is potentially translatable to high-field clinical MRI scanners for precision neuro-oncology imaging.

## Materials and Methods

### Animals and cell lines

All animal experiments were pre-approved by the competent institutional as well as national authorities, and carried out strictly adhering to European Directive 2010/63. A total of n=10 C57BL/6j male mice were used in this study, bred at the Champalimaud Foundation Vivarium, and housed with *ad libitum* access to food and water and 12h light cycles. GL261 mouse glioma cells were obtained from the Tumor Bank Repository at the National Cancer Institute (Frederick MD, USA). CT2A mouse glioma cells were kindly provided by Prof. Thomas Seyfried at Boston College (Boston MA, USA). Both cell lines were grown in RPMI-1640 culture medium supplemented with 2.0 g/l Sodium Bicarbonate, 0.285 g/l L-glutamine, 10% Fetal Bovine Serum (Gibco) and 1% Penicillin-Streptomycin solution. The cell lines tested negative for mycoplasma contamination using the IMPACT Mouse FELASA 1 test (Idexx-BioResearch, Ludwigsburg, Germany).

### Glioma models

Tumors were induced in previously described [66]. Briefly, intracranial stereotactic injection of 1 ×10^5^ GL261 or CT2A cells was performed in the caudate nucleus (n=5 and n=5 mice, respectively); analgesia (Meloxicam 1.0 mg/Kg s.c.) was administered 30 min before the procedure. Mice were anesthetized with isoflurane (1.5-2.0% in air) and immobilized on a stereotactic holder (Kopf Instruments, Tujunga/CA, USA) where they were warmed on a heating pad at 37 °C, while body temperature was monitored with a rectal probe (WPI ATC-2000, Hitchin, UK). The head was shaved with a small trimmer, cleaned with iodopovidone, and the skull exposed through an anterior-posterior incision in the midline with a scalpel. A 1 mm hole was drilled in the skull using a micro-driller, 0.1 mm posterior to the bregma and 2.32 mm lateral to the midline. The tumor cells (1×10^5^ in 4 μL PBS) were inoculated 2.35 mm below the cortical surface using a 10 µL Hamilton syringe (Hamilton, Reno NV, USA) connected to an automatic push-pull microinjector (WPI *Smartouch^TM^*, Sarasota FL, USA), by advancing the 26G needle 3.85 mm from the surface of the skull (∼1mm skull-to-brain surface distance), pulling it back 0.5 mm, and injecting at 2 μL/min rate. The syringe was gently removed 2 min after the injection had finished, the skin sutured with surgical thread (5/0 braided silk, Ethicon, San Lorenzo Puerto Rico) and wiped with iodopovidone. During recovery from anesthesia, animals were kept warm on a heating pad and given an opioid analgesic (Buprenorphine 0.05 mg/Kg s.c.) before returning to their cage. Meloxicam analgesia was repeatedly administered at 24- and 48-hours post-surgery.

### In vivo Studies

#### Longitudinal MRI

GBM-bearing mice were imaged every 5-7 days on a 1 Tesla Icon MRI scanner (Bruker BioSpin, Ettlingen, Germany; running *ParaVision 6.0.1* software), to measure tumor volumes. For this, each mouse was placed in the animal holder under anesthesia (1-2 % isoflurane in 31% O_2_), heated with a recirculating water blanket, and monitored for rectal temperature (36-37 °C) and breathing (60-90 BPM). Tumor volume was measured with T2-weighted ^1^H-MRI (*RARE* sequence, ×8 acceleration factor, repetition time TR = 2500 ms, echo time TE = 84 ms, 8 averages, 1 mm slice thickness, and 160×160 µm^2^ in-plane resolution), acquired in two orientations (coronal and axial). Each session lasted up to 30 min/animal.

#### End-point MRI and DMI

GBM-bearing mice with tumors ≥35 mm^3^ (longitudinal MRI assessment) were scanned on a 9.4T BioSpec MRI scanner (Bruker BioSpin, Ettlingen, Germany; running under *ParaVision 6.0.1*), using a ^2^H/^1^H transmit-receive surface coilset customized for the mouse brain (NeosBiotec, Pamplona, Spain), as described before [31]. Before each experiment, GBM-bearing mice fasted 4-6h, were weighed, and cannulated in the tail vein with a catheter connected to a home-built 3-way injection system filled with: 6,6′-^2^H_2_-glucose (1.6M in saline); Gd-DOTA (25 mM in saline); and with heparinized saline (10 U/mL). Mice were placed on the animal holder under anesthesia (as in 2.3.1). Coilset quality factors (Q) for ^1^H and ^2^H channels were estimated in the scanner for each sample based on the ratio of the resonance frequency (400.34 and 61.45 MHz, for protons and deuterium, respectively) to its bandwidth (full width at half-minimum of the wobbling curve during the initial tuning adjustments): 175±8 and 200±12, respectively. Mice were imaged first with T2-weighted ^1^H-MRI (*RARE* sequence, x8 acceleration factor, 3000 ms TR, 40 ms TE; 2 averages, 1 mm slice thickness, 70 µm in-plane resolution) in two orientations (coronal and axial). Then, the magnetic field homogeneity was optimized over the tumor region based on the water peak with ^1^H-MRS (*STEAM* localization: 6×6×3 mm volume of interest, i.e. 108 µL) using localized 1^st^ and 2^nd^ order shimming with the *MapShim Bruker* macro, leading to full widths at half-maximum (FWHM) of 28±5 Hz.

DMI was performed using a *slice-FID chemical-shift imaging* pulse sequence, with 175 ms TR, 256 spectral points sampled over a 1749 Hz window, and Shinnar-Le Roux RF pulse [67, 68] (0.42ms, 10kHz) with 55° flip angle, to excite a brain slice including the tumor: 18×18 mm field-of-view, and 2.27 mm slice thickness. After RF pulse calibration (using the natural abundance semi-heavy water peak, DHO), DGE-DMI data were acquired for 2h23min (768 repetitions), with i.v. bolus of 6,6′-^2^H_2_-glucose (2 mg/g, 4 µL/g injected over 30 s; Euroisotop, St Aubin Cedex, France). Data were sampled with an 8×8 matrix and 4-fold Fourier interpolated [69], rendering a 560 µm in-plane resolution. A reference T2-weighted image was additionally acquired with matching field-of-view and slice thickness, and 70 µm in-plane resolution.

Finally, animals underwent DCE T1-weighted ^1^H-MRI (*FLASH* sequence, 8° flip-angle, 16ms TR, 4 averages, 150 repetitions, 1 slice with 140 µm in-plane resolution and 2.27 mm thickness, FOV size and position matching the DGE-DMI experiment), with i.v. bolus injection of Gd-DOTA (0.1 mmol/Kg, injected over 30 s; Guerbet, Villepinte, France). Animals were then sacrificed, brains were removed, washed in PBS, and immersed in 4% PFA.

### MRI/DMI Processing

#### T2-weighted ^1^H-MRI

T2-weighted MRI data were processed in ImageJ 1.53a (Rasband, W.S., ImageJ, U. S. National Institutes of Health, Bethesda, Maryland, USA, https://imagej.nih.gov/ij/, 1997-2018). For each animal, the tumor region was manually delineated on each slice, and the sum of the areas multiplied by the slice thickness to estimate the volume, which was averaged across the two orientations acquired (coronal and axial).

#### DGE-DMI

DGE-DMI data were processed in MATLAB^®^ R2018b (Natick, Massachusetts: The MathWorks Inc.) and jMRUI 6.0b [70]. Each dataset was averaged to 12 min temporal resolution and noise regions outside the brain, as well as the olfactory bulb and cerebellum, were discarded, rendering a 4D spectral-spatial-temporal matrix of 256×32×32×12 points. After automated phase-correction of each spectrum, the 4D matrix was denoised with a tensor PCA denoising approach [32]. For this, a [8 8 8] window and tensor structure [1 2:3 4] were used for patch processing the spectral, spatial, and temporal dimensions with, whereas the *a priori* average standard deviation of the noise in each spectrum (calculated σ^2^) was used to avoid deleterious effects of spatially-correlated noise [34]. Then, these denoised spectra were analyzed voxel-wise by individual peak fitting with AMARES (similarly to the single-spectrum analysis reported previously in [52]), using a basis set for DHO (4.76 ppm: short- and long-T2 fractions [18]) and deuterium-labelled: glucose (Glc, 3.81 ppm), glutamate-glutamine (Glx, 2.36 ppm), and lactate (Lac, 1.31 ppm); relative linewidths referenced to the estimated short-T2 fraction of DHO, according to the respective T2 relaxation times reported by de Feyter *et al* [18]. The natural abundance DHO peak (DHOi) was further used to select and quantify both original and denoised spectra: SNR_DHOi_ >3.5 and 13.88 mM reference (assuming 80 % water content in the brain and 0.03 % natural abundance of DHO), respectively. Metabolite concentrations (CRLB<50%; otherwise discarded) were corrected for T1 and labeling-loss effects, according to the values reported by de Feyter *et al* (T1, ms: DHO, 320; Glc, 64; Glx, 146; Lac, 297) [18] and de Graaf *et al* (number of magnetically equivalent deuterons: DHO, 1; Glc, 2; Glx, 1.2; Lac, 1.7) [71], respectively. Thus, the concentration of each metabolite (m) at each time point was estimated as (Eq 1):

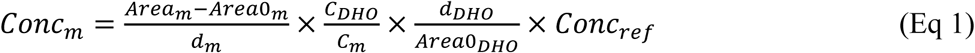

*Area* = peak area; *Area0* = average peak area before injection; *d* = number of magnetically equivalent deuterons corrected for labelling-loss effects; *C* = T1 correction factor (1-exp(-TR/T1)); and *Conc_ref_* = reference DHO concentration.

The time-course changes of ^2^H-labelled metabolite (Glc, Glx and Lac) concentrations were fitted using a modified version of the kinetic model reported by Kreis et al [19], to estimate the maximum rate of Glc consumption (total, *V_max_*) for Glx synthesis (mitochondrial oxidation, *V_glx_*) and Lac synthesis (glycolysis, *V_lac_*), and the confidence intervals for all estimated parameters:

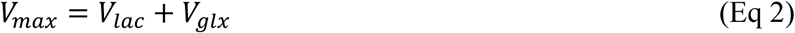

The coupled differential equations describing the concentration kinetics of each metabolite were:

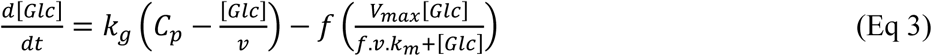

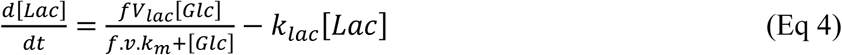

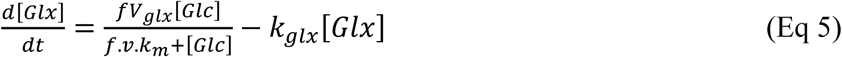

where: *k_g_*, apparent rate constant of glucose transfer between blood and tumor (min^−1^); *k_glx_*, apparent rate constant of Glx elimination (min^−1^); *k_lac_*, apparent rate constant of lactate elimination (min^−1^); *C*_*p*_ = *a*_1_ · *e*^−*kp*·*t*^, Glc concentration in plasma (mM); *a_1_*, the Glc concentration after the bolus injection (mM); and *k_p_*, the effective rate constant of labeled glucose transfer to tissue (min^−1^). As reported previously [31], the following parameters were fixed: fraction of deuterium enrichment (*f*), at 0.6 [19]; constant for glucose uptake (*k_m_*), at 10 mM [72, 73]; and the extravascular-extracellular volume fraction (*v*), at 0.22 – average estimation from DCE-T1-weighted MRI analysis (**Table S1**). All the other parameters were fitted without any restrictions to their range.

#### DCE T1-weighted MRI

DCE T1-weighted MRI data were processed with DCE@urLab [74], as before [31]. First, ROIs were manually delineated for each tumor and the time-course data was fitted with the Extended Tofts 2-compartment model [75], to derive the volume transfer constant between plasma and tumor extravascular-extracellular space (*k^trans^*), the washout rate between extravascular-extracellular space and plasma (*k_ep_*), and the extravascular-extracellular volume fraction (*v_e_*). Then, each dataset was reprocessed by down-sampling the original in-plane resolution to match the DGE-DMI experiment (0.56×0.56×2.27 mm^3^), and fitting the time-course data pixel-wise with the Extended Tofts 2-compartment model to derive *k^trans^*, *k_ep_*, and *v_e_* maps (pixels with root-mean square error >0.005 discarded).

### Histopathology and Immunohistochemistry

Whole brains fixed in 4% PFA were embedded in paraffin and sectioned at 30 different levels on the horizontal plane, spanning the whole tumor area. 4 µm sections were stained with H&E (Sigma-Aldrich, St. Louis MO, USA), digitized (Nanozoomer, Hamamatsu, Japan), and analyzed by an experimental pathologist blinded to experimental groups, according to previously established criteria [31]. Then, QuPath v0.4.3 built-in tools [76] were used to highlight different tumor regions: Tumor ROIs, corresponding to the bulk tumor, were delineated first with “create threshold” and then manually corrected; P-Margin ROIs, including areas of peritumoral infiltration, were delineated with “expand annotations” by expanding 100 µm the tumor margin toward the adjacent brain parenchyma; Infiltrative ROIs, corresponding to specific infiltrative regions, were manually annotated. Between 3 to 6 sections of each tumor were also immunostained for Ki67 (mouse anti-ki67, BD, San Jose CA, USA; blocking reagent, M.O.M ImmPRESS kit, Vector Laboratories, Burlingame CA, USA; liquid DAB^+^, Dako North America Inc, Carpinteria CA, USA), digitized (Nanozoomer, Hamamatsu, Japan), and analyzed with QuPath built-in tools [76] for Tumor and P-Margin ROIs, defined as detailed above. Thus, Ki67^+/-^ cells were counted semi-automatically to determine the total number of cells, the cell density, and the proliferation index (% Ki67^+^ cells) as the average across slices for each ROI, and respective Tumor/P-Margin ratios. This procedure was repeated for each animal. In addition, one histologic section corresponding to each DGE-DMI slice was immunostained for Iba-1 (rabbit anti-Iba-1, Fujifilm Wako PCC, Osaka, Japan; NovolinkTM Polymer, Leica Biosystems, UK; liquid DAB+, Dako North America Inc, Carpinteria CA, USA), digitalized (Philips UFS v1.8.6614 slide scanner) and analyzed in QuPath. Tumor region and peritumoral margin regions were automatically annotated as outlined above, and Iba-1 positive staining was quantified across all annotations using the threshold tools, adjusted for each slide to account for variations in staining intensity, to calculate the percentage of Iba-1 positive area: (Iba-1+ area / total annotation area) * 100.

### Statistical analyses

Data were analyzed in MATLAB^®^ R2018b (Natick, Massachusetts: The MathWorks Inc.) using the two-tailed Student’s *t-*test, either unpaired (comparing different animal cohorts) or paired (comparing the same animal cohort in different conditions). Differences at the 95% confidence level (p=0.05) were considered statistically significant. Correlation analyses were carried out with the Pearson R coefficient. Error bars indicate standard deviation unless indicated otherwise.

## Acknowledgments

This work was supported by: H2020-MSCA-IF-2018, ref. 844776 (RVS); FCT CEEC-IND4ed, ref 2021.02777.CEECIND/CP1675/CT0003 (RNH); and the Champalimaud Foundation. The authors thank Dr. Thomas Seyfried for access to the CT2A cell line and helpful discussion, and the Vivarium of the Champalimaud Centre for the Unknown, a research infrastructure of CONGENTO co-financed by Lisbon Regional Operational Programme (*Lisboa2020*), under the PORTUGAL 2020 Partnership Agreement, through the European Regional Development Fund (ERDF) and *Fundação para a Ciência e Tecnologia* (Portugal), under the project LISBOA-01-0145-FEDER-022170.

## Appendix A. Supplementary Data

**SimoesRV_SupplementaryData.pdf**

