## Supplemental Data for "Deuterium Metabolic Imaging Phenotypes Mouse Glioblastoma Heterogeneity Through Glucose Turnover Kinetics"

| | |
|---|---|
| Supplementary Figures (S1 to S12) | 2 |
| Supplementary Tables (S1 to S3) | 15 |

### Supplementary Figures

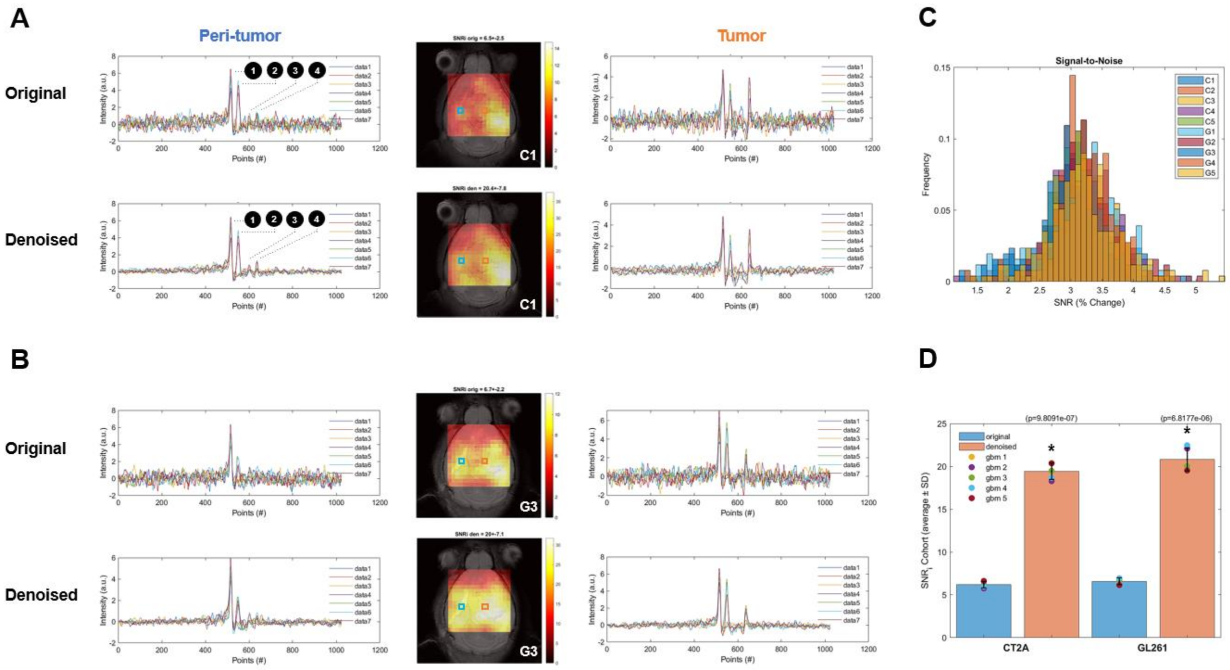


**Figure S1 - Tensor PCA denoising improves DGE-DMI SNR in mouse GBM.** Examples of CT2A (C1, **A**) and GL261 (G3, **B**) subjects, showing the SNR maps (center) from the original data and after tensor PCA denoising, as well as examples of time-course spectra from tumor (right-side) and peri-tumoral regions (left-side) in each condition – voxel positions overlaid on the SNR maps (orange and blue, respectively). **C** Histograms of pixel-wise SNR fold-changes after tensor PCA denoising for each subject (color-coded). **D.** Significant SNR increase in CT2A and GL261 cohorts after tensor PCA denoising. **1**, semi-heavy water signal (DHO); **2**, 6,6′ -$^{2}H_2$-glucose (Glc); **3**, 4,4′ -$^{2}$H-glutamate-glutamine (Glx); **4**, 3,3′ -$^{2}$H-lactate (Lac). * p<0.001.

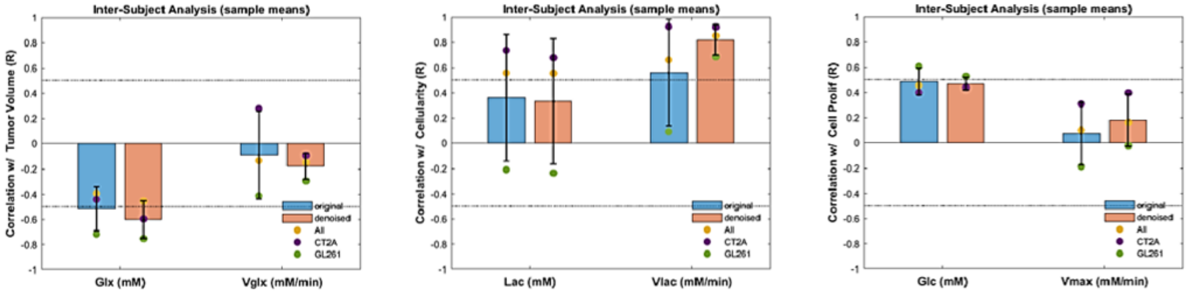


**Figure S2 - Gross inter-subject correlations of averaged metabolic maps.** Correlation coefficients (Pearson R) are displayed for CT2A (purple), GL261 (green), and pooled cohorts (CT2A+GL261, yellow), contrasting results from original data (blue) and tensor PCA denoised data (orange). **Left-side**, glutamate-glutamine accumulation (Glx) and glucose consumption rate for its synthesis ($V_{glx}$) vs tumor volume. **Center**, lactate accumulation (Lac) and glucose consumption rate for its synthesis ($V_{lac}$) vs cellularity. **Right-side**, glucose accumulation (Glc) and maximum consumption rate ($V_{max}$) vs cell proliferation.

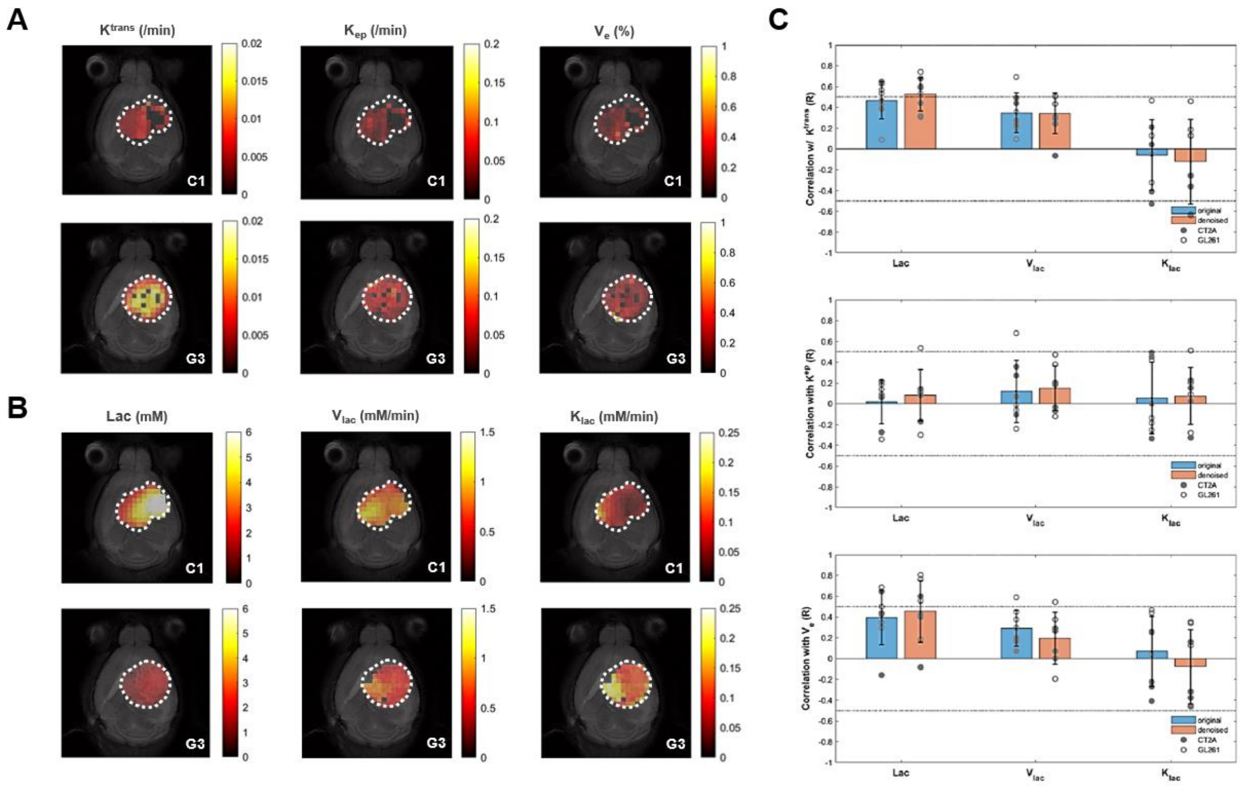


**Figure S3 - Intra-tumor pixel-wise correlations between metabolic and permeability metrics**. Examples of CT2A (C1) and GL261 (G3) subjects, displaying DCE-T1 permeability maps (**A**) and glucose-derived lactate maps derived from DGE-DMI after tensor PCA denoising (**B**). **C** Correlations coefficients (Pearson R) displayed for each tumor in CT2A and GL261 cohorts (closed and open circles, respectively), contrasting results from original data (blue) and tensor PCA denoised data (orange).

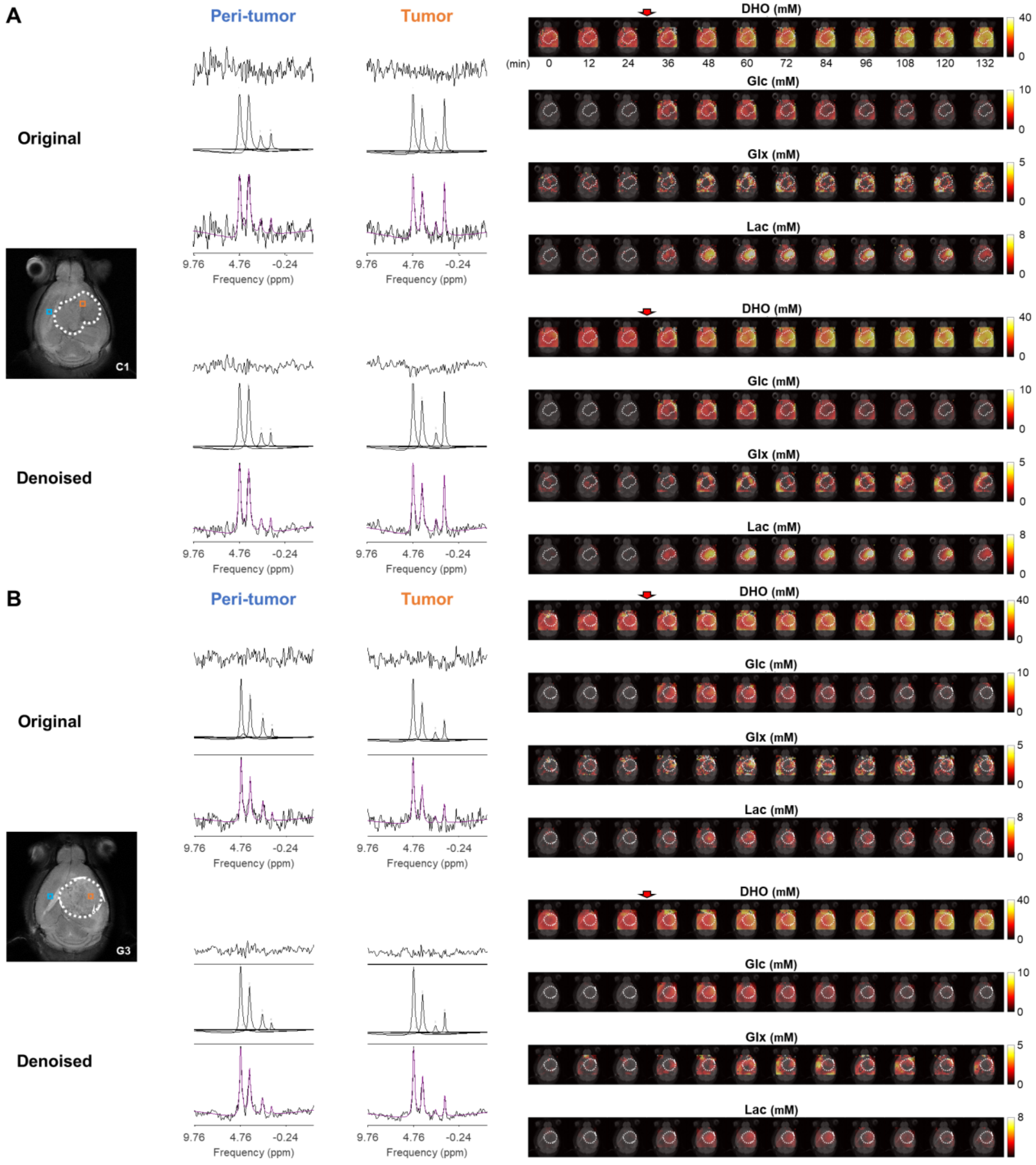


**Figure S4 - Quantification of DGE-DMI data.** Examples of CT2A (C1, **A**) and GL261 (G3, **B**) subjects, and respective tumor regions (dashed lines), showing: on left-side, the improved spectral quality and respective quantification in tumor and peritumor regions (bottom, raw spectrum with overlaid estimation in purple; center, individual components; top, residual); and on the right-side, time-course metabolic concentration maps of (top-to-bottom) DHO, Glc, Glx, and Lac, following Glc i.v. injection.

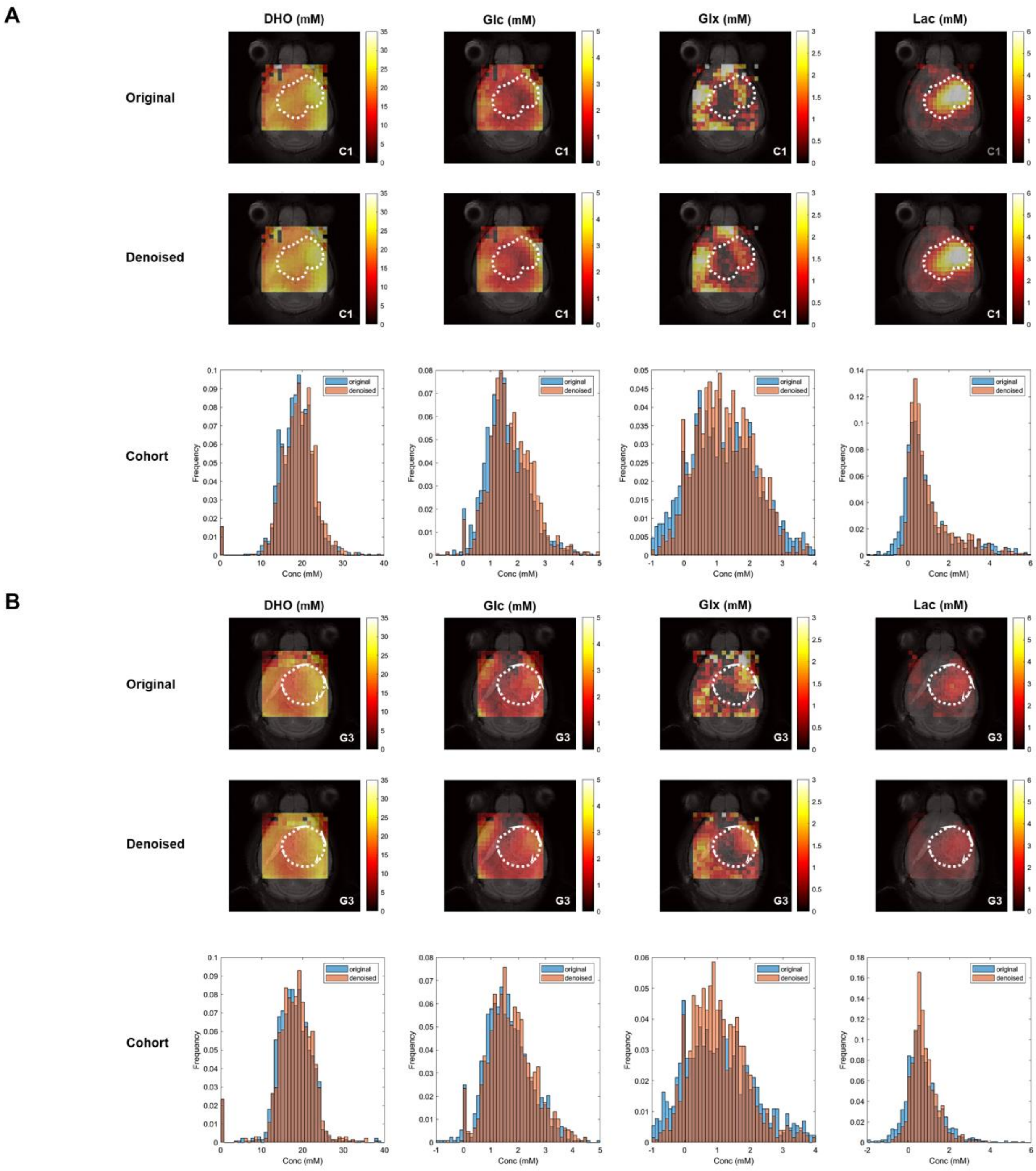


**Figure S5 - Tensor PCA denoising has no overall effect on pixel distributions of DGE-DMI metabolic concentration maps in pooled GBM cohorts**. CT2A (**A**) and GL261 (**B**) cohorts, showing examples (subjects C1 and G3, respectively) of *de novo* concentration maps generated from original data (first row) and tensor PCA denoised data (second row): DHO, glucose (Glc), glucose-derived glutamine-glutamate (Glx) and lactate (Lac) (top, left-to-right). Total pixel distributions of the total cohorts are also displayed for each map (bottom).

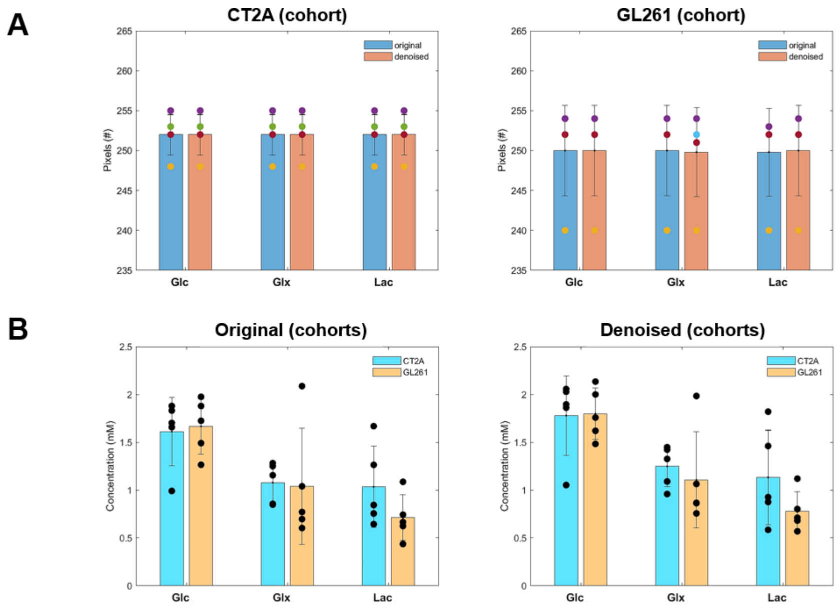


**Figure S6 - Tensor PCA denoising has no effect on pixel detectability or GBM cohort differences of DGE-DMI time-course average metabolic concentration maps.** Time-course average of pixel detected in CT2A (left-side) and GL261 (right-side) cohorts, comparing tensor PCA denoising vs original. **B** Cohort differences of original (left-side) and tensor PCA denoised data (right-side), comparing CT2A vs GL261. No significant differences detected.

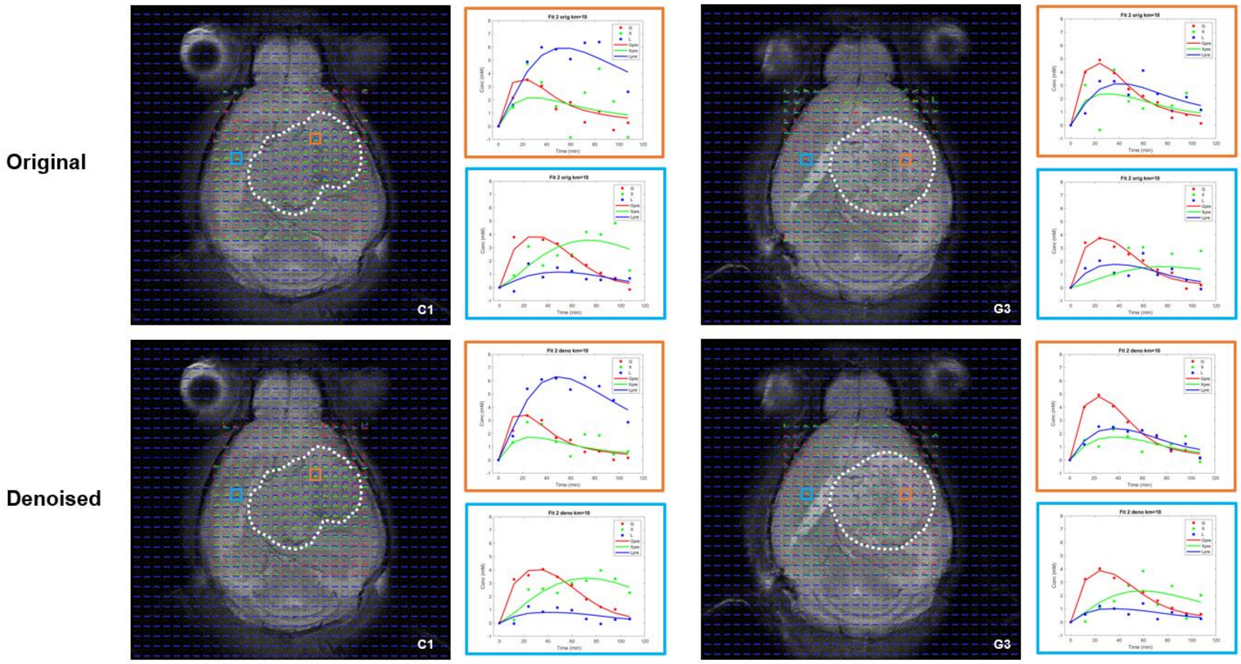


**Figure S7 - Kinetic modeling of DGE-DMI time-course concentration maps data.** Examples of CT2A (C1, **left-side**) and GL261 (G3, **right-side**) subjects and respective tumor regions (dashed lines), with overlaid time-course concentration plots for each metabolite – Glc (red), Glx (green), and Lac (blue) – and respective kinetic fitting (straight lines, same color codes). Voxels from tumor (orange) and peritumoral (light blue) regions are shown enlarged, displaying original data (**top**) and tensor PCA denoised data (**bottom**).

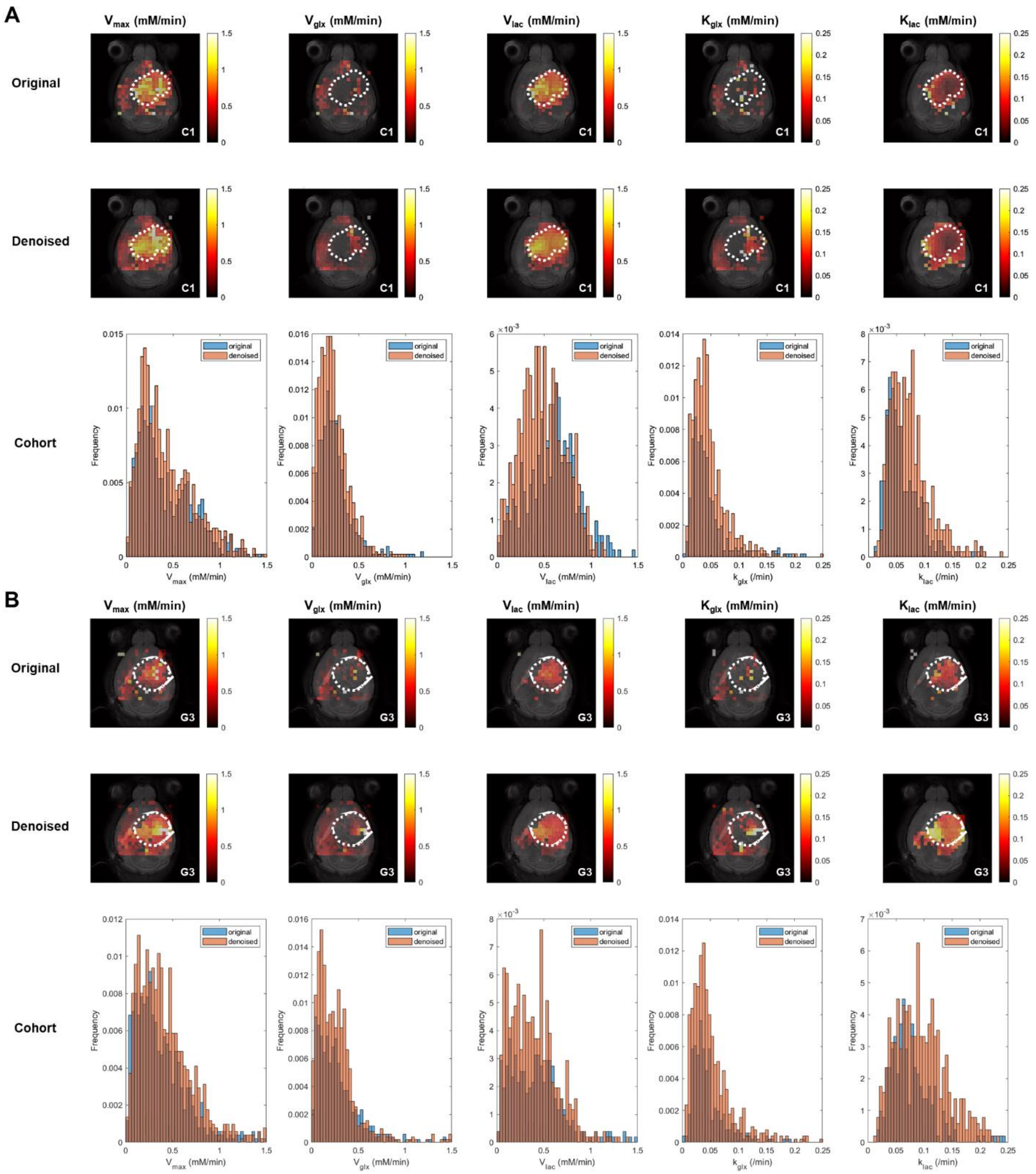


**Figure S8 - Tensor PCA denoising increases pixel densities of DGE-DMI metabolic flux maps in pooled GBM cohorts.** CT2A (**A**) and GL261 (**B**) cohorts, showing examples (subjects C1 and G3, respectively) of glucose flux maps (top) generated from original data (first row) and tensor PCA denoised data (second row): maximum consumption rate ($V_{max}$) and respective consumption rates for lactate synthesis ($V_{lac}$) and elimination ($k_{lac}$), and glutamate-glutamine synthesis ($V_{glx}$) and elimination ($k_{glx}$). Total pixel distributions of the total cohorts are also displayed for each map (bottom).

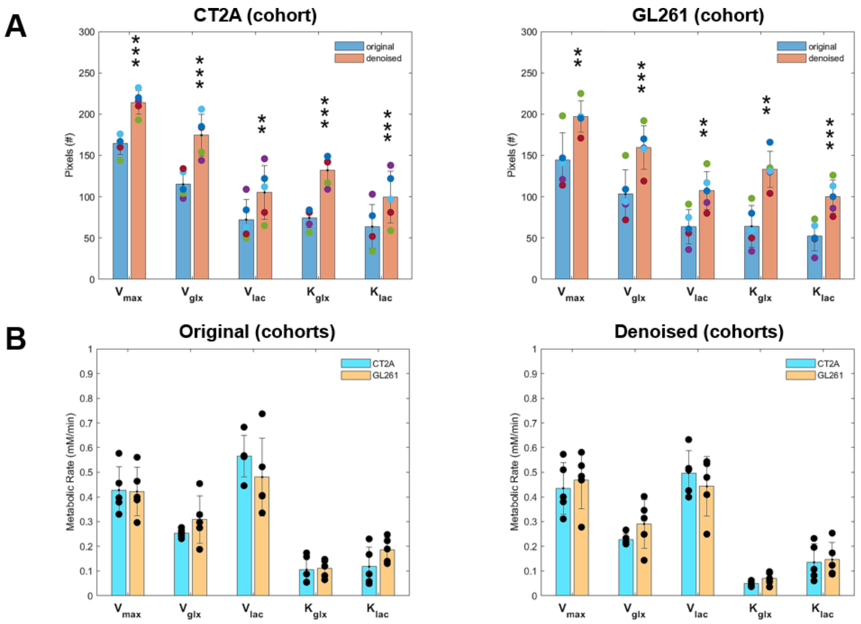


**Figure S9 - Tensor PCA denoising improves pixel detectability without affecting GBM cohort differences of DGE-DMI metabolic flux maps. A** Time-course average of pixels detected in CT2A (left-side) and GL261 (right-side) cohorts, comparing tensor PCA denoising vs original (** $p<0.01$, *** $p<0.001$) – overall pixel detectability: CT2A, +53±18%; GL261, +73±30%. **B** Cohort differences of original (left-side) and tensor PCA denoised data (right-side), comparing CT2A vs GL261: no significant differences detected.

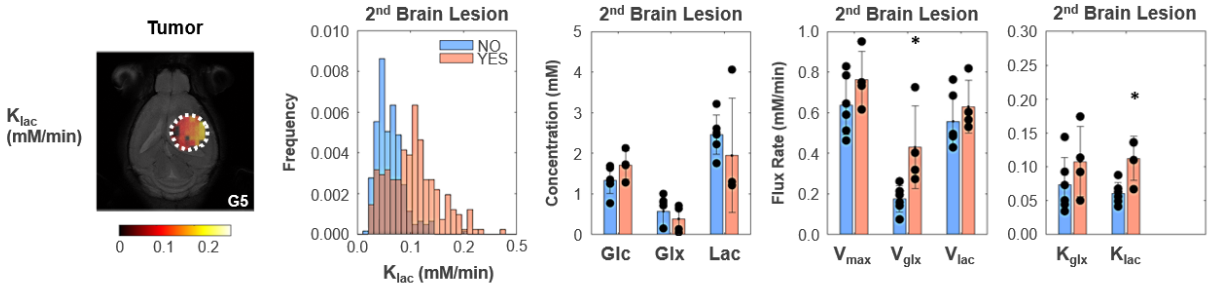


**Figure S10 - Tumor metabolic changes mirror GBM infiltration and migration leading to secondary brain lesions.** Metabolic map (**left-side:** C5 tumor), histogram distributions (**center:** pooled GL261 and CT2A cohorts), and group comparison of mean values (**right-side:** bar plots), indicating significantly higher rates of glutamate-glutamine synthesis and lactate consumption/elimination in primary tumors displaying secondary brain lesions. Secondary lesion, with (n=4) vs without (n=6): * $p<0.05$ ($k_{lac}$ +84%, p=0.010; and $V_{glx}$ +146%, p=0.019); unpaired *t*-test. Error bars: standard deviation. elimination

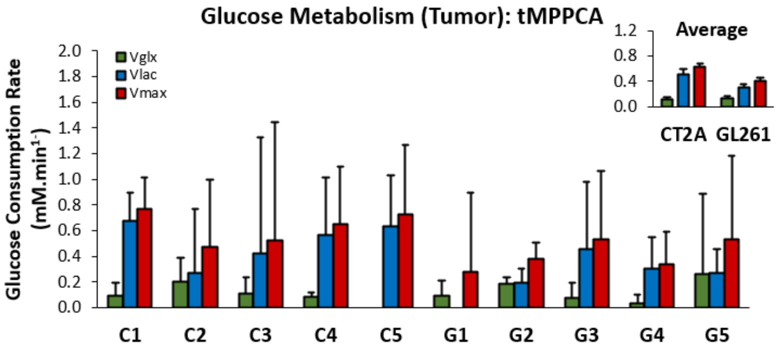


**Figure S11 - Glucose consumption rates in mouse GBM tumors following ROI averaged raw data**. FID averaging within each tumor ROI (CT2A, C1-5; and GL261, G1-5) was followed by Fourier Transform, tensor PCA spectral denoising, spectral quantification, and kinetic modeling, to derive the metrics displayed: synthesis rates of glucose-derived glutamate-glutamine ($V_{glx}$, green) and lactate ($V_{lac}$, blue), and maximum consumption rate of glucose ($V_{max}$, red). Upper-right box displaying cohort averages. Plots: estimates ± SE (mM/min).

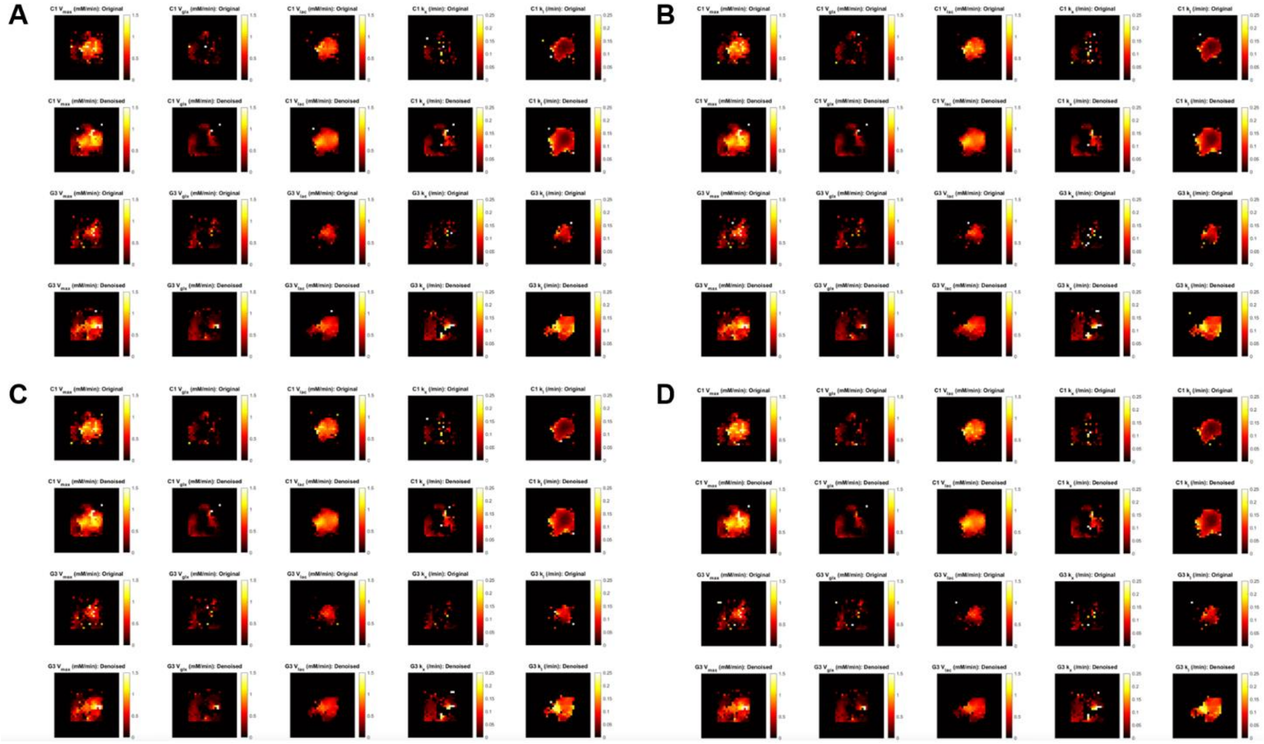


**Figure S12 - Effect of changing $v_e$ on the metabolic maps derived from kinetic modeling.** Fixing the extracellular volume fraction to: **A**, cohort minima (CT2A, 0.14; GL261, 0.21); **B**, cohort means (CT2A, 0.18; GL261, 0.26); **C**, average of A-C values (pooled cohorts, 0.19). **D**, pooled cohort average (0.22). Each panel displays CT2A (C1, rows 1-2) and GL261 (G3, rows 3-4) subjects, showing results from original data (rows 1 and 3) and tensor PCA denoised data (rows 2 and 4) for the metabolic maps (left-to-right): maximum glucose consumption rate ($V_{max}$), glucose consumption for synthesis of glutamate-glutamine ($V_{glx}$) and lactate ($V_{lac}$), and their respective consumption rates ($k_{glx}$ and $k_{lac}$).

### Supplementary Tables

**Table S1. Tumor cohorts.** Animal information and metrics obtained from multi-modal *in vivo* MRI and *post-mortem* histopathology and immunostaining.

| | Parameter | CT2A TUMORS C1 | C2 | C3 | C4 | C5 | CT2A COHORT (mean±SE) | GL261 TUMORS G1 | G2 | G3 | G4 | G5 | GL261 COHORT (mean±SE) | GL261 vs CT2A (%) | (p) [c] | POOLED COHORTS (mean±SE) |
|---|---|---|---|---|---|---|---|---|---|---|---|---|---|---|---|---|
| Animal | Batch | 1 | 2 | 2 | 2 | 2 | N.A. | 1 | 1 | 2 | 2 | 2 | N.A. | N.A. | N.A. | N.A. |
| | Gender | male | male | male | male | male | N.A. | male | male | male | male | male | N.A. | N.A. | N.A. | N.A. |
| | **Days PI** | 18 | 45 | 22 | 28 | 37 | 30 ± 5 | 18 | 18 | 16 | 17 | 17 | 17 ± 0 | **42.7%** | **0.03215** | 24 ± 3 |
| | **Weight (g)** | 24.3 | 23.0 | 26.6 | 26.4 | 28.3 | 25.7 ± 0.9 | 22.1 | 24.5 | 21.9 | 20.8 | 22.7 | 22.4 ± 0.6 | **-12.9%** | **0.01741** | 24.1 ± 0.8 |
| T2w MRI | Tumor Volume ($mm^3$) | 113.9 | 41.2 | 52.3 | 34.7 | 59.8 | 60.4 ± 14.1 | 41.0 | 49.1 | 74.6 | 64.6 | 54.1 | 56.7 ± 5.9 | -6.2% | 0.81283 | 58.5 ± 7.2 |
| DGE-DMI | WL (Hz) | 31.2 | 33.3 | 24.7 | 20.4 | 36.5 | 29.2 ± 2.9 | 20.4 | 22.6 | 27.9 | 29 | 30.1 | 26.0 ± 1.9 | -11.0% | 0.38367 | 27.6 ± 1.7 |
| | SNRi orig (average) | 6.5 | 5.7 | 6.1 | 6.0 | 6.6 | 6.2 ± 0.2 | 6.2 | 6.9 | 6.7 | 6.9 | 6.1 | 6.6 ± 0.2 | 6.1% | 0.15013 | 6.4 ± 0.1 |
| | SNRi den (average) | 20.4 | 18.3 | 19.5 | 18.7 | 20.4 | 19.5 ± 0.4 | 20.0 | 22.1 | 20.0 | 22.5 | 19.5 | 20.8 ± 0.6 | 7.0% | 0.10721 | 20.1 ± 0.4 |
| | ***Tumor (ROI):*** | | | | | | | | | | | | | | | |
| | Glc (mM) [a] | 1.73 | 0.77 | 1.33 | 1.25 | 1.35 | 1.29 ± 0.15 | 1.69 | 1.64 | 1.71 | 2.13 | 1.28 | 1.69 ± 0.13 | 31.4% | 0.08249 | 1.49 ± 0.12 |
| | Glx (mM) [a] | 0.71 | 0.04 | 0.82 | 0.79 | 0.72 | 0.62 ± 0.15 | 1.00 | 0.15 | 0.64 | 0.05 | 0.13 | 0.40 ± 0.18 | -35.5% | 0.37892 | 0.51 ± 0.12 |
| | **Lac (mM)** [a] | 4.06 | 2.22 | 2.46 | 2.61 | 3.22 | 2.91 ± 0.33 | 2.50 | 1.76 | 1.22 | 1.30 | 1.21 | 1.60 ± 0.25 | **-45.2%** | **0.01283** | 2.26 ± 0.29 |
| | $V_{max}$(mM.min$^{-1}$) [b] | 0.951 | 0.512 | 0.645 | 0.590 | 0.828 | 0.705 ± 0.080 | 0.784 | 0.463 | 0.733 | 0.616 | 0.750 | 0.669 ± 0.059 | -5.1% | 0.72736 | 0.687 ± 0.047 |
| | $V_{glx}$ (mM.min$^{-1}$) [b] | 0.273 | 0.159 | 0.195 | 0.142 | 0.262 | 0.207 ± 0.027 | 0.214 | 0.075 | 0.318 | 0.726 | 0.403 | 0.347 ± 0.109 | 68.1% | 0.24636 | 0.277 ± 0.058 |
| | $V_{lac}$ (mM.min$^{-1}$) [b] | 0.818 | 0.484 | 0.488 | 0.495 | 0.685 | 0.594 ± 0.068 | 0.764 | 0.429 | 0.604 | 0.565 | 0.530 | 0.578 ± 0.055 | -2.6% | 0.86151 | 0.586 ± 0.041 |
| | $K_{glx}$ (mM.min$^{-1}$) [b] | 0.114 | 0.051 | 0.073 | 0.035 | 0.093 | 0.073 ± 0.014 | 0.045 | 0.144 | 0.093 | 0.254 | 0.174 | 0.142 ± 0.036 | 94.4% | 0.11066 | 0.107 ± 0.021 |
| | **$K_{lac}$ (mM.min$^{-1}$)** [b] | 0.067 | 0.042 | 0.056 | 0.049 | 0.063 | 0.055 ± 0.005 | 0.088 | 0.068 | 0.136 | 0.136 | 0.110 | 0.108 ± 0.013 | **94.3%** | **0.00619** | 0.082 ± 0.011 |
| | ***Peritumoral Rim (ROI):*** | | | | | | | | | | | | | | | |
| | Glc (mM) [a] | 2.227 | 0.898 | 1.699 | 1.785 | 1.737 | 1.67 ± 0.22 | 1.633 | 1.325 | 1.503 | 1.958 | 1.461 | 1.58 ± 0.11 | -5.6% | 0.70857 | 1.62 ± 0.11 |
| | Glx (mM) [a] | 0.551 | 0.382 | 0.846 | 1.026 | 1.023 | 0.77 ± 0.13 | 1.477 | 0.472 | 0.812 | 0.561 | 0.698 | 0.80 ± 0.18 | 5.0% | 0.86554 | 0.78 ± 0.10 |
| | **Lac (mM)** [a] | 2.005 | 1.096 | 1.476 | 1.144 | 1.586 | 1.46 ± 0.17 | 1.272 | 0.976 | 0.755 | 0.871 | 0.840 | 0.94 ± 0.09 | **-35.5%** | **0.02466** | 1.20 ± 0.12 |
| | $V_{max}$(mM.min$^{-1}$) [b] | 0.597 | 0.427 | 0.440 | 0.528 | 0.536 | 0.505 ± 0.032 | 0.621 | 0.286 | 0.426 | 0.717 | 0.746 | 0.559 ± 0.088 | 10.6% | 0.58343 | 0.532 ± 0.045 |
| | $V_{glx}$ (mM.min$^{-1}$) [b] | 0.125 | 0.161 | 0.207 | 0.219 | 0.216 | 0.186 ± 0.018 | 0.306 | 0.060 | 0.171 | 0.326 | 0.611 | 0.295 ± 0.093 | 58.8% | 0.28113 | 0.240 ± 0.048 |
| | $V_{lac}$ (mM.min$^{-1}$) [b] | 0.587 | 0.398 | 0.355 | 0.577 | 0.475 | 0.479 ± 0.047 | 0.557 | 0.240 | 0.391 | 0.548 | 0.313 | 0.410 ± 0.063 | -14.4% | 0.40435 | 0.444 ± 0.039 |
| | $K_{glx}$ (mM.min$^{-1}$) [b] | 0.056 | 0.062 | 0.046 | 0.049 | 0.060 | 0.055 ± 0.003 | 0.038 | 0.027 | 0.051 | 0.114 | 0.187 | 0.083 ± 0.030 | 52.4% | 0.36967 | 0.069 ± 0.015 |
| | $K_{lac}$ (mM.min$^{-1}$) [b] | 0.091 | 0.071 | 0.077 | 0.141 | 0.089 | 0.094 ± 0.012 | 0.109 | 0.060 | 0.120 | 0.240 | 0.094 | 0.124 ± 0.030 | 32.7% | 0.37862 | 0.109 ± 0.016 |
| DCE-T1w MRI | ***Tumor (ROI):*** | | | | | | | | | | | | | | | |
| | **$K^{trans}$ (min$^{-1}$)** | 0.00527 | 0.00288 | N.A. | 0.00465 | 0.00445 | 0.00431 ± 0.00051 | 0.00532 | 0.00947 | 0.00976 | N.A. | 0.00959 | 0.00854 ± 0.00107 | **97.9%** | **0.01199** | 0.00642 ± 0.00097 |
| | $K^{ep}$ (min$^{-1}$) | 0.03455 | 0.03037 | N.A. | 0.03920 | 0.03059 | 0.03368 ± 0.00208 | 0.02128 | 0.03598 | 0.05167 | N.A. | 0.04561 | 0.03864 ± 0.00663 | 14.7% | 0.50174 | 0.03616 ± 0.00335 |
| | v (%) | 0.192 | 0.140 | N.A. | 0.148 | 0.230 | 0.178 ± 0.021 | 0.346 | 0.272 | 0.211 | N.A. | 0.227 | 0.264 ± 0.030 | 48.7% | 0.05658 | 0.221 ± 0.024 |
| | ***Peritumoral Rim (ROI):*** | | | | | | | | | | | | | | | |
| | $K^{trans}$ (min$^{-1}$) | 0.00294 | 0.00109 | N.A. | 0.00190 | 0.00084 | 0.00169 ± 0.00047 | 0.00116 | 0.00257 | 0.00240 | N.A. | 0.00230 | 0.00211 ± 0.00032 | 24.5% | 0.49496 | 0.00190 ± 0.00028 |
| | $K^{ep}$ (min$^{-1}$) | 0.04049 | 0.04568 | N.A. | 0.01839 | 0.03472 | 0.03482 ± 0.00592 | 0.02515 | 0.05097 | 0.08493 | N.A. | 0.04581 | 0.05171 ± 0.01240 | 48.5% | 0.26486 | 0.04327 ± 0.00712 |
| | v (%) | 0.085 | 0.026 | N.A. | 0.150 | 0.067 | 0.082 ± 0.026 | 0.105 | 0.106 | 0.026 | N.A. | 0.103 | 0.085 ± 0.020 | 3.3% | 0.93572 | 0.083 ± 0.015 |
| Histology | Phenotype (H&E score) | I | III | III | I | III | 0/5 [d] | IV | IV | IV | IV | IV | 5/5 [d] | N.A. | N.A. | N.A. |
| | Infiltration (+ yes; – no) | – | – | – | – | – | 0/5 [e] | + | + | + | + | + | 5/5 [e] | N.A. | N.A. | N.A. |
| | Distant Migration (+ yes; – no) | + | – | – | – | – | 1/5 [e] | – | – | + | + | + | 3/5 [e] | N.A. | N.A. | N.A. |
| | ***Tumor (ROI):*** | | | | | | | | | | | | | | | |
| | Area ($10^7$ μm$^2$) | 1.70 | 0.84 | 0.80 | 1.02 | 1.15 | 1.10 ± 0.16 | 1.11 | 1.24 | 1.59 | 4.01 | 1.15 | 1.82 ± 0.56 | 65.1% | 0.24978 | 1.46 ± 0.30 |
| | Cellularity ($10^5$ cells) | 1.20 | 0.67 | 0.69 | 0.94 | 0.96 | 0.89 ± 0.10 | 0.56 | 0.60 | 0.85 | 0.66 | 0.47 | 0.63 ± 0.06 | -29.8% | 0.05224 | 0.76 ± 0.07 |
| | **Cell Density ($10^{-3}$ cells/um$^2$)** | 7.07 | 8.10 | 8.24 | 9.21 | 8.45 | 8.22 ± 0.34 | 5.06 | 4.81 | 5.35 | 5.17 | 4.07 | 4.89 ± 0.22 | **-40.4%** | **0.00004** | 6.56 ± 0.59 |
| | Cell Proliferation (Ki67$^+$ %) | 52.2 | 65.0 | 61.7 | 72.8 | 68.7 | 64.1 ± 3.5 | 76.8 | 77.8 | 76.1 | 70.0 | 62.1 | 72.5 ± 2.9 | 13.2% | 0.10187 | 68.3 ± 2.6 |
| | Mgl/Mp infiltration (% area) | 2.8 | N.A: | 4.7 | 7.9 | 1.4 | 4.2 ± 1.4 | 3.0 | 0.6 | 0.1 | 2.6 | 2.2 | 1.7 ± 0.6 | -58.9% | 0.12320 | 2.8 ± 0.8 |
| | ***Peritumoral Rim (ROI):*** | | | | | | | | | | | | | | | |
| | Area ($10^7$ μm$^2$) | 0.45 | 0.33 | 0.25 | 0.36 | 0.34 | 0.35 ± 0.03 | 0.33 | 0.32 | 0.30 | 0.31 | 0.31 | 0.31 ± 0.01 | -9.3% | 0.35993 | 0.33 ± 0.02 |
| | Cellularity ($10^5$ cells) | 0.08 | 0.07 | 0.04 | 0.06 | 0.06 | 0.06 ± 0.01 | 0.05 | 0.05 | 0.06 | 0.17 | 0.06 | 0.08 ± 0.02 | 20.9% | 0.59409 | 0.07 ± 0.01 |
| | Cell Density ($10^{-3}$ cells/um$^2$) | 1.87 | 2.04 | 1.77 | 1.84 | 1.64 | 1.83 ± 0.07 | 1.55 | 1.48 | 2.03 | 6.91 | 1.89 | 2.77 ± 1.04 | 51.4% | 0.39236 | 2.30 ± 0.52 |
| | Cell Proliferation (Ki67$^+$ %) | 16.6 | 27.8 | 25.5 | 24.3 | 17.4 | 22.3 ± 2.3 | 18.4 | 20.5 | 16.7 | 18.8 | 18.2 | 18.5 ± 0.6 | -17.0% | 0.14291 | 20.4 ± 1.3 |
| | **Mgl/Mp infiltration (% area)** | 8.7 | N.A. | 15.7 | 10.9 | 4.8 | 10.0 ± 2.3 | 4.5 | 1.5 | 0.2 | 1.6 | 2.9 | 2.1 ± 0.7 | **-78.6%** | **0.00839** | 5.6 ± 1.7 |

$K^{trans}$, volume transfer constant between plasma and tumor extravascular-extracellular space; $k_{ep}$, washout rate between extravascular-extracellular space and plasma; $v_e$, extravascular-extracellular volume fraction; WL, water linewidth at half-maximum of the water peak; *Vglx*, maximum rate of Glc consumption for Glx synthesis (mM. min−1); *Vlac*, maximum rate of Glc consumption for Lac synthesis (mM. min−1); *Vmax*, maximum rate of total Glc consumption (mM. min−1); [a] Temporal average (post-injection); [b] Kinetic flux rate; [c] 2-tailed, unpaired t-Test (highlighted p<0.05); [d] tumors with score IV (Supplementary Table 2); [e] tumors with infiltration, migration, or both.

**Table S2. Histopathologic evaluation.** Analysis of H&E histologic sections from each tumor and scoring according to phenotypic features of their stromal-vascular fraction.

| SAMPLE | TUMOR | | | | | | | | VESSELS | | | BRAIN | | | RESULTS |
|---|---|---|---|---|---|---|---|---|---|---|---|---|---|---|---|
| ID | primary location | secondary location | growth pattern[a] | tumor cell morphology | cell-cell interaction | stroma | tumor border[a] | tumor necrosis | density | instability | mimicry | vascular invasion[b] | intercallation | brain damage | Phenotype score[c] |
| C1 | cerebral nuclei: caudoputamen | midbrain | expansile | large, poligonal to spindle | cohesive | non-cystic, non-oedematous, non-hemorrhagic | outward pushing | minimal | moderate | mild | not evident | not evident | 0 or minimal | minimal: compressive, ischemic | I |
| C2 | cerebral nuclei: caudoputamen | none | expansile | large, poligonal to spindle | poorly cohesive | cystic, oedematous, hemorrhagic | outward pushing | moderate | moderate | marked | present | not evident | mild | mild: compressive, ischemic | III |
| C3 | cerebral nuclei: caudoputamen | none | expansile | large, poligonal to spindle | cohesive | non-cystic, non-oedematous, poorly hemorrhagic | outward pushing | mild | moderate | moderate | present | not evident | 0 or minimal | minimal: compressive, ischemic | III |
| C4 | cerebral nuclei: caudoputamen | none | expansile | large, poligonal to spindle | cohesive | non-cystic, non-oedematous, non-hemorrhagic | outward pushing | minimal | moderate | mild | not evident | not evident | 0 or minimal | minimal: compressive, ischemic | I |
| C5 | cerebral nuclei: caudoputamen | none | expansile | large, spindle | poorly cohesive | cystic, oedematous, hemorrhagic | outward pushing | moderate | moderate | marked | present | not evident | mild | mild: compressive, hemorrhagic | III |
| G1 | cerebral nuclei: caudoputamen | none | expansile, infiltrative | large, poligonal, multinucleated | poorly cohesive | cystic, oedematous, non-hemorrhagic | outward pushing and solid strand collective migration | minimal | low | mild | present | not evident | moderate | minimal: compressive | IV |
| G2 | cerebral nuclei: caudoputamen | none | expansile, infiltrative | large, poligonal, multinucleated | poorly cohesive | cystic, oedematous, non-hemorrhagic | outward pushing and solid strand collective migration | minimal | low | mild | present | not evident | moderate | minimal: compressive | IV |
| G3 | cerebral nuclei: caudoputamen | infiltrating ipsiateral and contralateral lateral ventricle | expansile | large, poligonal, multinucleated | poorly cohesive at 1ary tumor | cystic, oedematous, non-hemorrhagic | outward pushing | minimal | moderate | moderate | present | not evident | 0 or minimal | minimal: compressive, ischemic | IV |
| G4 | cerebral nuclei: caudoputamen | infiltrating ipsiateral and contralateral lateral ventricle | expansile | large, poligonal, multinucleated | non-cohesive | cystic, oedematous, hemorrhagic | outward pushing and solid strand collective migration | minimal | moderate | marked | present | not evident | 0 or minimal | minimal: compressive, ischemic | IV |
| G5 | cerebral nuclei: caudoputamen | infiltrating ipsiateral lateral ventricle | expansile | large, poligonal, multinucleated | non-cohesive | cystic, oedematous, hemorrhagic | outward pushing and solid strand collective migration | minimal | moderate | marked | present | not evident | 0 or minimal | minimal: compressive, ischemic | IV |

a predominant; b 0 (absent) or 1 (present); c progression phase; d tumors studied *in vivo* under acute hypoxia.

**Table S3. Tumor-to-border ratios in CT2A and GL261 cohorts.** Animal information and metrics obtained from multi-modal *in vivo* MRI and *post-mortem* histopathology and immunostaining. Initial values taken from Supplementary table 1.

| | | CT2A TUMORS | | | | | CT2A COHORT | GL261 TUMORS | | | | | GL261 COHORT | CT2A vs GL261 | | POOLED COHORTS |
|---|---|---|---|---|---|---|---|---|---|---|---|---|---|---|---|---|
| | Parameter | C1 | C2 | C3 | C4 | C5 | (mean±SE) | G1 | G2 | G3 | G4 | G5 | (mean±SE) | (%) | (p) [c] | (mean±SE) |
| DGE-DMI | ***Tumor/PT-Rim (ROI):*** | | | | | | | | | | | | | | | |
| | **Glc** [a] | 0.78 | 0.86 | 0.78 | 0.70 | 0.78 | 0.78 ± 0.02 | 1.03 | 1.24 | 1.14 | 1.09 | 0.88 | 1.08 ± 0.06 | **37.9%** | **0.00185** | 0.93 ± 0.06 |
| | Glx [a] | 1.29 | 0.10 | 0.98 | 0.77 | 0.70 | 0.77 ± 0.20 | 0.68 | 0.32 | 0.79 | 0.10 | 0.19 | 0.42 ± 0.14 | -45.8% | 0.17812 | 0.59 ± 0.13 |
| | **Lac** [a] | 2.03 | 2.03 | 1.67 | 2.28 | 2.03 | 2.01 ± 0.10 | 1.97 | 1.80 | 1.61 | 1.49 | 1.44 | 1.66 ± 0.10 | **-17.1%** | **0.03814** | 1.83 ± 0.09 |
| | $V_{max}$ [b] | 1.59 | 1.20 | 1.47 | 1.12 | 1.54 | 1.38 ± 0.10 | 1.26 | 1.62 | 1.72 | 0.86 | 1.01 | 1.29 ± 0.17 | -6.5% | 0.65134 | 1.340 ± 0.092 |
| | $V_{glx}$ [b] | 2.18 | 0.99 | 0.94 | 0.65 | 1.21 | 1.20 ± 0.26 | 0.70 | 1.24 | 1.86 | 2.22 | 0.66 | 1.34 ± 0.31 | 11.9% | 0.73566 | 1.266 ± 0.193 |
| | $V_{lac}$ [b] | 1.39 | 1.22 | 1.38 | 0.86 | 1.44 | 1.26 ± 0.11 | 1.37 | 1.79 | 1.54 | 1.03 | 1.70 | 1.49 ± 0.13 | 18.3% | 0.21603 | 1.371 ± 0.089 |
| | $K_{glx}$ [b] | 2.03 | 0.81 | 1.60 | 0.70 | 1.54 | 1.34 ± 0.25 | 1.18 | 5.38 | 1.81 | 2.22 | 0.93 | 2.31 ± 0.80 | 72.4% | 0.28244 | 1.821 ± 0.428 |
| | **$K_{lac}$** [b] | 0.74 | 0.59 | 0.73 | 0.35 | 0.71 | 0.62 ± 0.07 | 0.80 | 1.13 | 1.14 | 0.57 | 1.17 | 0.96 ± 0.12 | **54.9%** | **0.04011** | 0.792 ± 0.087 |
| DCE-T1w MRI | ***Tumor/PT-Rim (ROI):*** | | | | | | | | | | | | | | | |
| | $K^{trans}$ | 1.79254 | 2.64653 | N.A. | 2.45347 | 5.27242 | 3.04124 ± 0.76587 | 4.58606 | 3.67981 | 4.06757 | N.A. | 4.17236 | 4.12645 ± 0.18626 | 35.7% | 0.21773 | 3.58385 ± 0.41855 |
| | $K^{ep}$ | 0.85324 | 0.66484 | N.A. | 2.13101 | 0.88103 | 1.13253 ± 0.33627 | 0.84638 | 0.70592 | 0.60833 | N.A. | 0.99583 | 0.78911 ± 0.08447 | -30.3% | 0.36019 | 0.96082 ± 0.17312 |
| | v | 2.25528 | 5.42510 | N.A. | 0.98829 | 3.41765 | 3.022 ± 0.942 | 3.29250 | 2.58069 | 8.17438 | N.A. | 2.20769 | 4.064 ± 1.389 | 34.5% | 0.55739 | 3.543 ± 0.801 |
| Histology | ***Tumor/PT-Rim (ROI):*** | | | | | | | | | | | | | | | |
| | Area | 3.76 | 2.56 | 3.23 | 2.81 | 3.37 | 3.15 ± 0.21 | 3.31 | 3.89 | 5.32 | 12.78 | 3.77 | 5.81 ± 1.77 | 84.8% | 0.17363 | 4.48 ± 0.95 |
| | Cellularity | 14.17 | 9.84 | 15.42 | 14.41 | 17.28 | 14.22 ± 1.22 | 11.07 | 12.61 | 13.98 | 3.89 | 8.13 | 9.93 ± 1.80 | -30.2% | 0.08391 | 12.08 ± 1.25 |
| | **Cell Density** | 3.78 | 3.97 | 4.66 | 5.00 | 5.16 | 4.51 ± 0.27 | 3.26 | 3.25 | 2.63 | 0.75 | 2.15 | 2.41 ± 0.46 | **-46.6%** | **0.00453** | 3.46 ± 0.43 |
| | **Cell Proliferation** | 3.2 | 2.3 | 2.4 | 3.0 | 3.9 | 3.0 ± 0.3 | 4.2 | 3.8 | 4.6 | 3.7 | 3.4 | 3.9 ± 0.2 | **32.3%** | **0.02568** | 3.5 ± 0.2 |
| | Mgl/Mp infiltration (% area) | 0.3 | N.A. | 0.3 | 0.7 | 0.3 | 0.4 ± 0.1 | 0.7 | 0.4 | 0.5 | 1.6 | 0.8 | 0.8 ± 0.2 | 96.6% | 0.18177 | 0.6 ± 0.1 |

$K^{trans}$, volume transfer constant between plasma and tumor extravascular-extracellular space; $k_{ep}$, washout rate between extravascular-extracellular space and plasma; $v_e$, extravascular-extracellular volume fraction; WL, water linewidth at half-maximum of the water peak; *Vglx*, maximum rate of Glc consumption for Glx synthesis (mM. min−1); *Vlac*, maximum rate of Glc consumption for Lac synthesis (mM. min−1); *Vmax*, maximum rate of total Glc consumption (mM. min−1); [a] Temporal average (post-injection); [b] Kinetic flux rate; [c] 2-tailed, unpaired t-Test (highlighted $p<0.05$).
